# Alpha phase-amplitude tradeoffs predict visual perception

**DOI:** 10.1101/2021.05.25.445552

**Authors:** Camille Fakche, Rufin VanRullen, Philippe Marque, Laura Dugué

## Abstract

Spontaneous alpha oscillations (~10 Hz) have been associated with various cognitive functions, including perception. Their phase and amplitude independently predict cortical excitability and subsequent perceptual performance. Yet, the causal role of alpha phase-amplitude tradeoffs on visual perception remains ill-defined. We aimed to fill this gap and tested two clear predictions from the Pulsed Inhibition theory according to which alpha oscillations are associated with periodic functional inhibition. (1) High alpha amplitude induces cortical inhibition at specific phases, associated with low perceptual performance, while at opposite phases, inhibition decreases (potentially increasing excitation) and perceptual performance increases. (2) Low alpha amplitude is less susceptible to these phasic (periodic) pulses of inhibition, leading to overall higher perceptual performance. Here, cortical excitability was assessed in humans using phosphene (illusory) perception induced by single pulses of transcranial magnetic stimulation (TMS) applied over visual cortex at perceptual threshold, and its post-pulse evoked activity recorded with simultaneous electroencephalography (EEG). We observed that pre-pulse alpha phase modulates the probability to perceive a phosphene, predominantly for high alpha amplitude, with a non-optimal phase for phosphene perception between −π/2 and −π/4. The pre-pulse non-optimal phase further leads to an increase in post-pulse evoked activity (ERP), in phosphene-perceived trials specifically. Together, these results show that alpha oscillations create periodic inhibitory moments when alpha amplitude is high, leading to periodic decrease of perceptual performance. This study provides strong causal evidence in favor of the Pulsed Inhibition theory.

**Visual Abstract:** 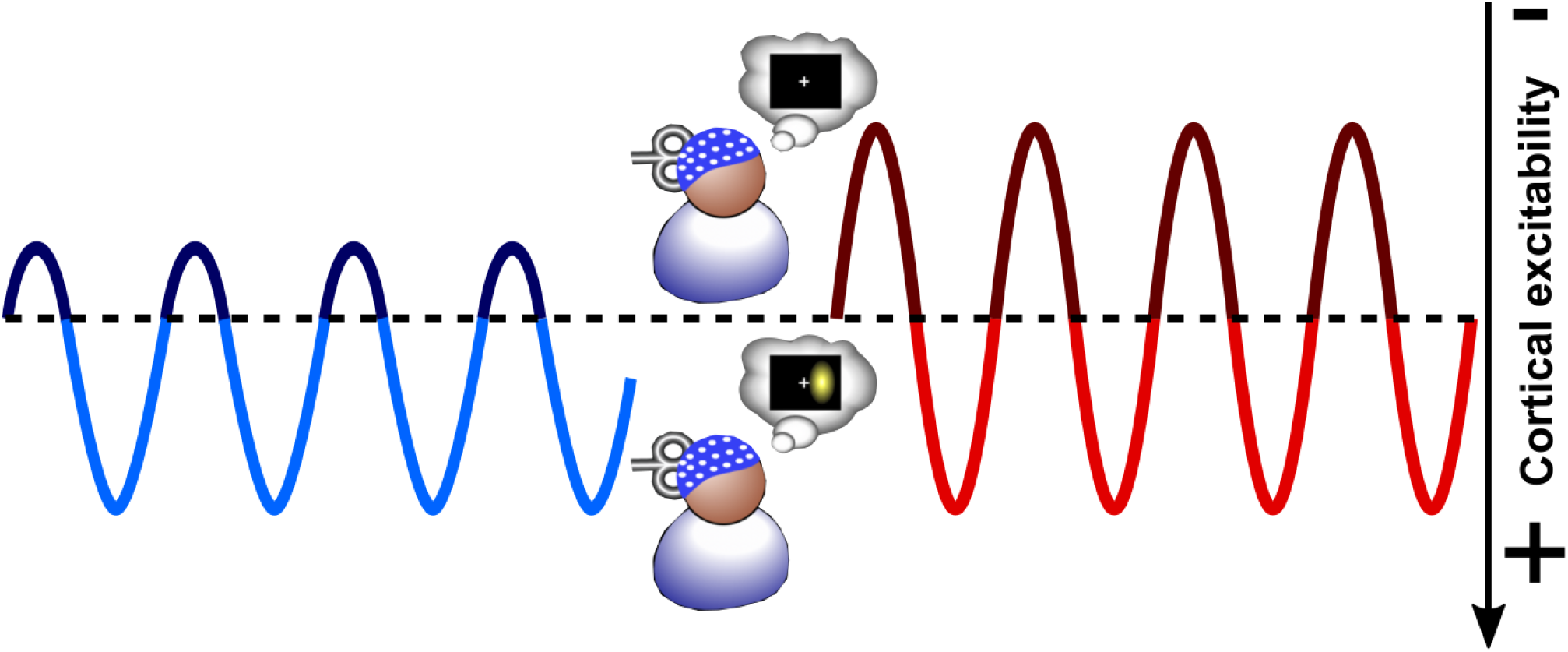

**Significance Statement:** The Pulsed Inhibition theory predicts that the functional inhibition induced by high alpha oscillations’ amplitude is periodic, with specific phases decreasing neural firing and perceptual performance. In turn, low alpha oscillations’ amplitude is less susceptible to phasic moments of pulsed inhibition leading to overall higher perceptual performance. Using TMS with simultaneous EEG recordings in humans, we found that specific phases of spontaneous alpha oscillations (~10 Hz) decrease cortical excitability and the subsequent perceptual outcomes predominantly when alpha amplitude is high. Our results provide strong causal evidence in favor of the Pulsed Inhibition theory.

## Introduction

Alpha brain oscillations (8-12 Hz) play a role in various cognitive functions, including visual perception (VanRullen, 2016a; Dugué and VanRullen, 2017; Clayton et al., 2018; Samaha et al., 2020; Kienitz et al., 2021), and are proposed to support active functional inhibition (Pfurtscheller and Lopes da Silva, 1999). Specifically, low parieto-occipital alpha amplitude recorded in human is correlated with a higher probability to perceive a near-threshold visual stimulus (Ergenoglu et al., 2004; Händel et al., 2011; Sauseng et al., 2005; Thut et al., 2006; van Dijk et al., 2008). Similarly, specific alpha phases lead to better detection performance while opposite phases lead to impaired performance (Varela et al., 1981; Busch et al., 2009; Mathewson et al., 2009; Dugué et al., 2011a; Samaha et al., 2015). Perception fluctuates periodically over time, along with the phase of alpha oscillations.

Alpha oscillations’ amplitude and phase further seem to predict cortical excitability. Spiking activity recorded in macaques is higher at the trough of the alpha cycle, and for lower alpha amplitude (Bollimunta et al., 2008; Haegens et al., 2015, 2011; van Kerkoerle et al., 2014). Moreover, functional Magnetic Resonance Imaging studies have shown that the cortical blood oxygenation level-dependent response fluctuates along with the phase of alpha oscillations in visual cortex (V1/V2; Scheeringa et al., 2011), and increases when alpha amplitude decreases (Goldman et al., 2002; Moosmann et al., 2003). Critically, transcranial magnetic stimulation (TMS) studies went beyond such correlational evidence. When applied over V1/V2, single-pulse TMS can elicit phosphenes (illusory percepts) depending on cortical state, i.e., when cortical excitability is sufficiently high. Phosphene perception leads to a higher event-related potential (ERP) than the absence of percept (Dugué et al., 2011a; Taylor et al., 2010; Samaha et al., 2017). Interestingly, studies have shown that phosphene perception is higher for low pre-pulse alpha amplitude (Romei et al., 2008; Samaha et al., 2017), and fluctuates according to alpha phase (Dugué et al., 2011a; Samaha et al., 2017). These studies independently suggest that both alpha oscillations’ amplitude and phase modulate cortical excitability and causally predict visual perception. Yet, their joint causal effects are still ill-defined.

Here, we address this question in the framework of the Pulsed Inhibition theory (Jensen and Mazaheri, 2010; Klimesch et al., 2007; Mathewson et al., 2011), which makes two clear predictions regarding alpha phase-amplitude tradeoffs: (1) high alpha amplitude exhibits states of cortical inhibition at specific phases associated with lower perceptual performance (non-optimal phases); while 2) low alpha amplitude is less susceptible to phasic pulsed inhibition, leading to high perceptual performance. A few studies have investigated the phase-amplitude tradeoffs (i.e., effect of the interaction between phase and amplitude) of low frequency oscillations on sensory perception and motor functions in human (**Table 1**). Although most of these studies observed a phase effect on task performance exclusively (or stronger) for high-alpha (or lower frequencies) amplitude (Ai and Ro, 2014; Alexander et al., 2020; Bonnefond and Jensen, 2015; Herrmann et al., 2016; Hussain et al., 2019; Kizuk and Mathewson, 2017; Mathewson et al., 2009; Ng et al., 2012; Spitzer et al., 2016), one found a phase effect for low-alpha amplitude exclusively (Busch and VanRullen, 2010); some found no difference between high- and low-alpha amplitude (Harris et al., 2018; Milton and Pleydell-Pearce, 2016); and finally, some found no phase effect (Madsen et al., 2019; Zoefel and Heil, 2013). Importantly, most of these studies are correlational (**Table 1**). Only two (Hussain et al., 2019; Madsen et al., 2019) investigated the causal link between spontaneous alpha oscillations’ amplitude and phase, cortico-spinal excitability –assessed with TMS to evoke a motor-evoked potential (MEP) on the hand muscle– and the subsequent motor performance. Other studies found a causal phase effect of spontaneous alpha oscillations on cortico-spinal excitability and the associated MEP for high-alpha amplitude, but used the absence of alpha oscillations as control thus not comparing phase-effects between a high- and low-alpha amplitude condition (Bergmann et al., 2019; Schaworonkow et al., 2019, 2018; Stefanou et al., 2018; Zrenner et al., 2018).

**Table 1.**
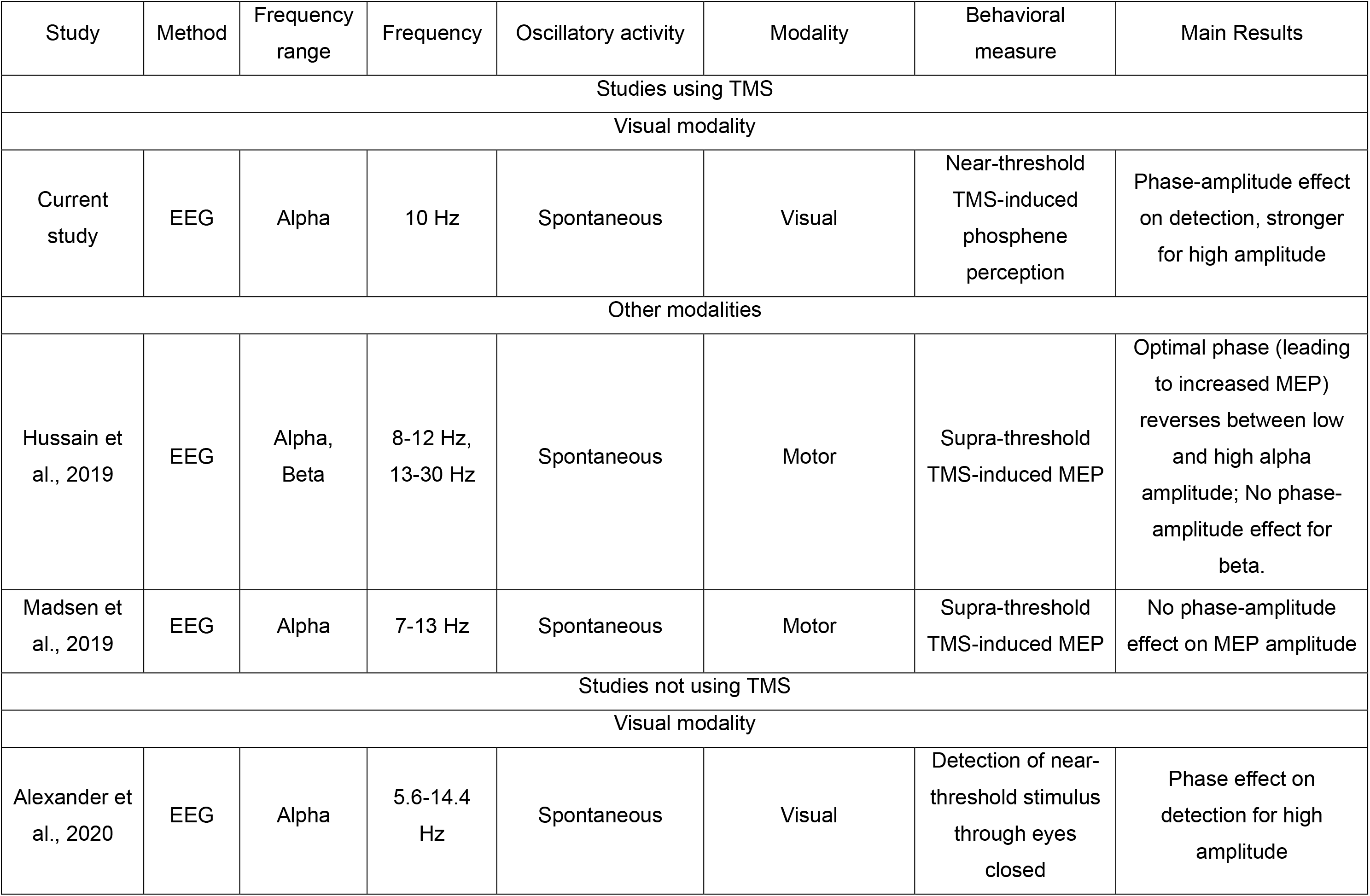
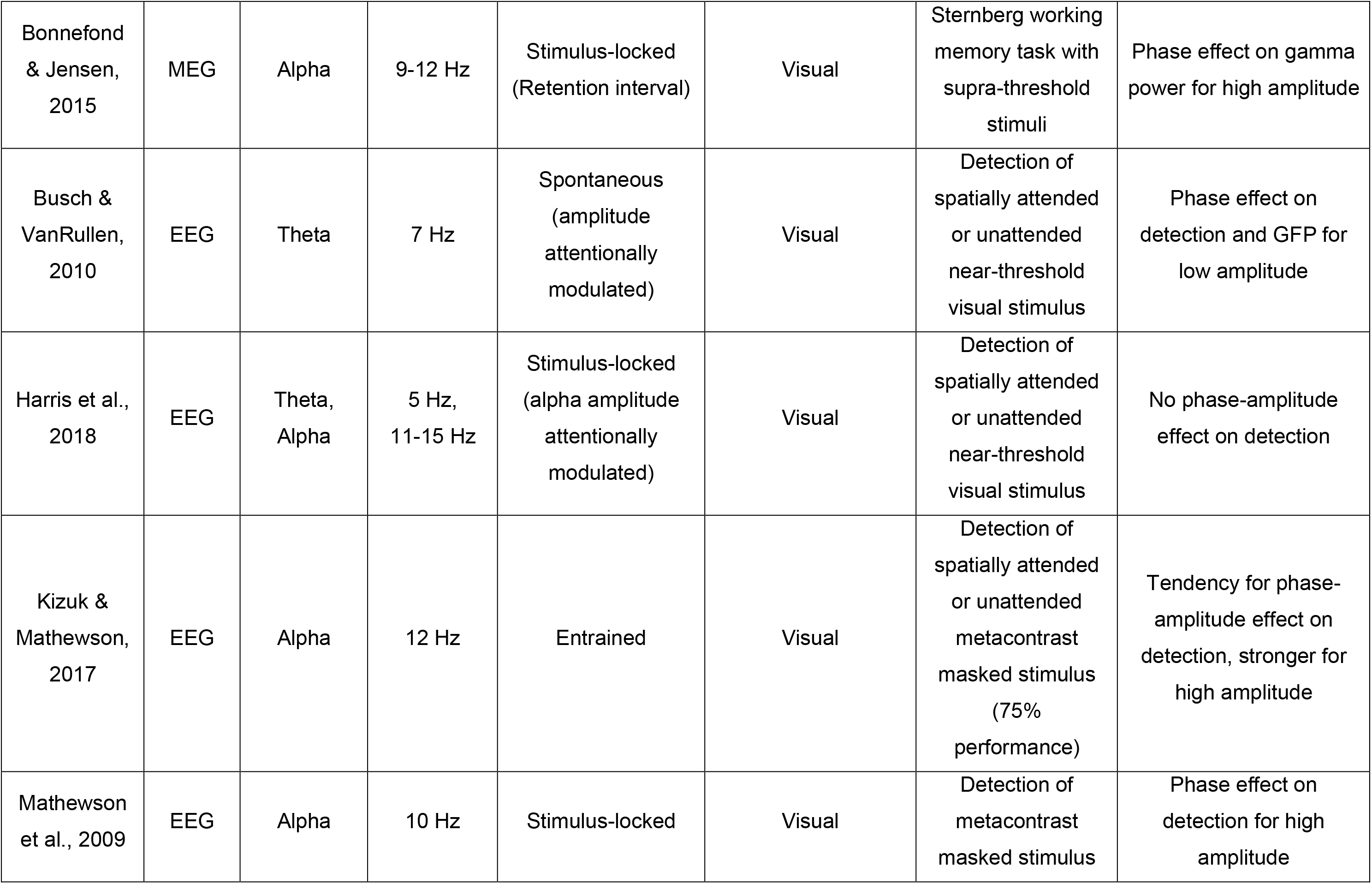
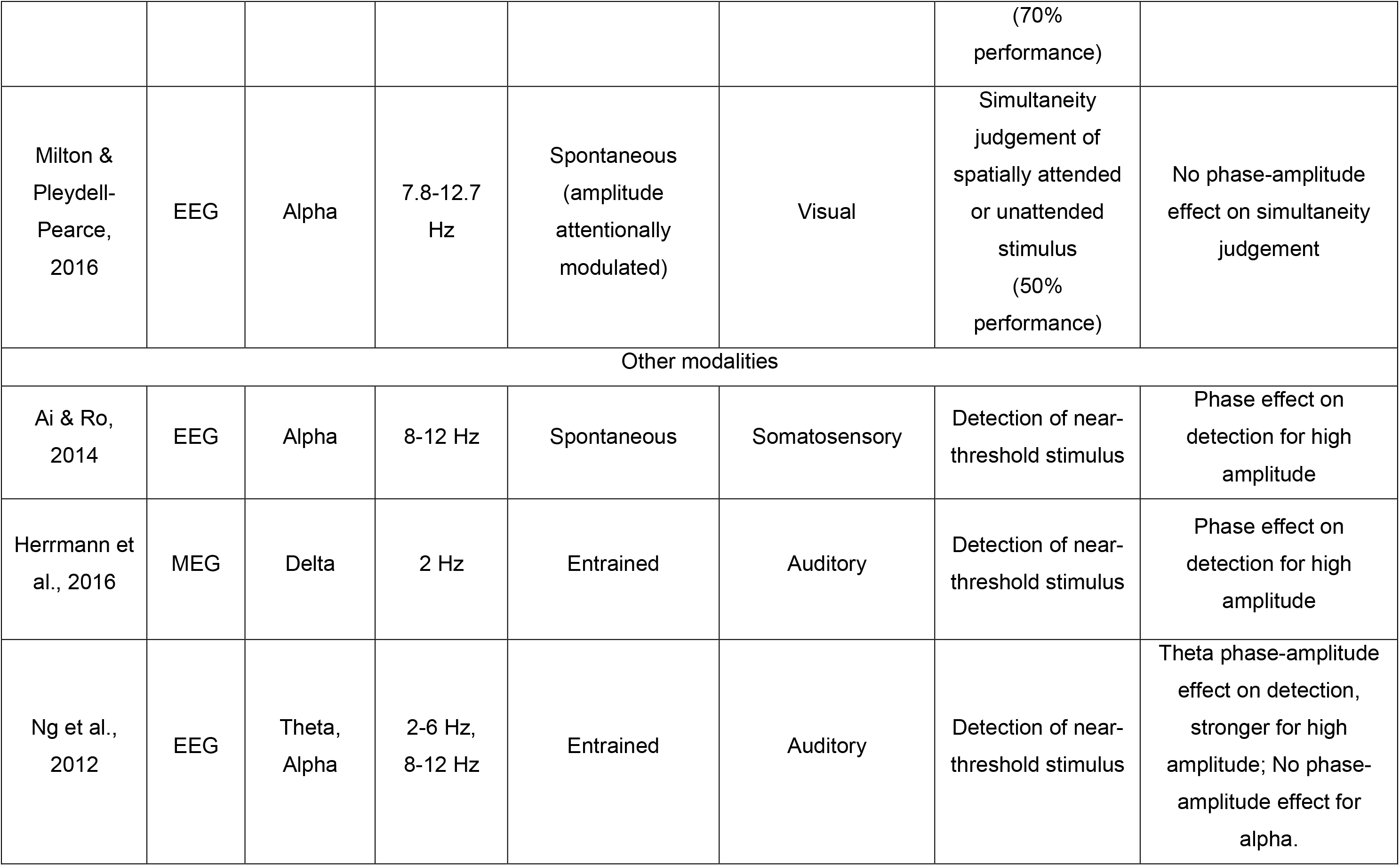
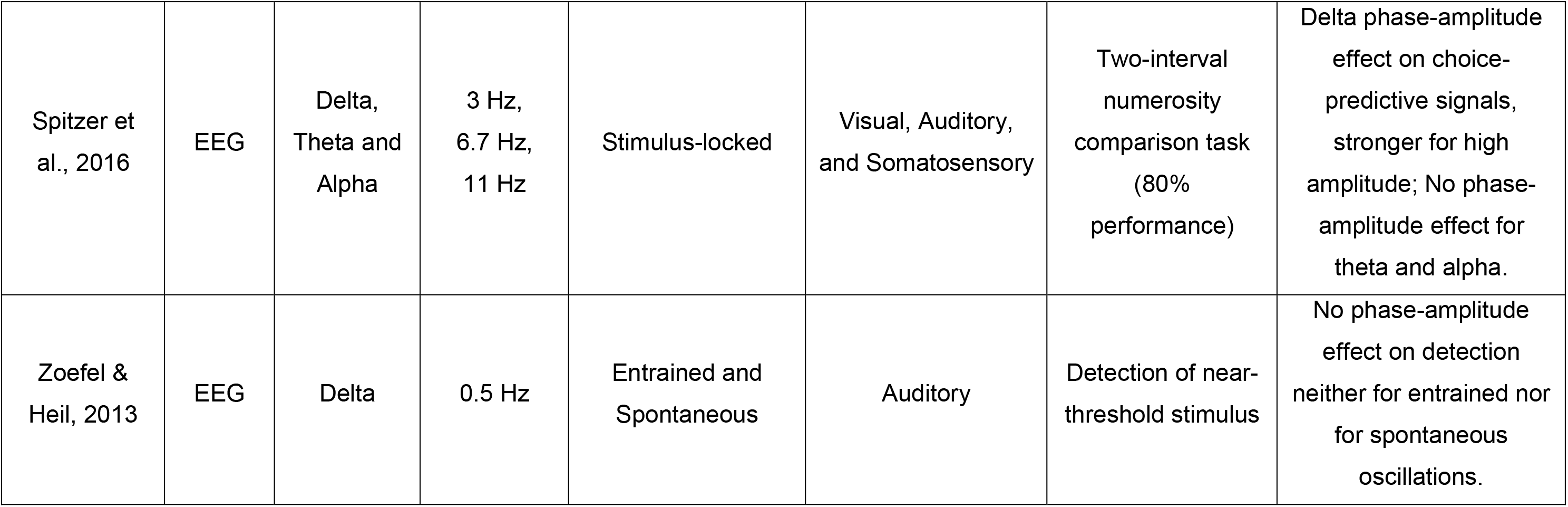
Electro/Magneto-encephalography (EEG/MEG) experiments investigating oscillations phase-amplitude tradeoffs on behavioral performance in human. Note that studies independently investigating the phase and the amplitude of oscillations were not included in this table. We only selected studies specifically investigating the interaction between the instantaneous phase and the amplitude on behavioral performance. Entrained oscillations: oscillations with a non-random phase distribution induced by a repetitive stimulus presentation. Stimulus-locked oscillations: oscillations with a non-random phase distribution induced by a single stimulus. Spontaneous oscillations: oscillations with a random phase distribution. GFP, Global Field Power. MEP, Motor Evoked Potential. TMS, Transcranial Magnetic Stimulation.

| Study | Method | Frequency range | Frequency | Oscillatory activity | Modality | Behavioral measure | Main Results |
| --- | --- | --- | --- | --- | --- | --- | --- |
| Studies using TMS |  |  |  |  |  |  |  |
| Visual modality |  |  |  |  |  |  |  |
| Current study | EEG | Alpha | 10 Hz | Spontaneous | Visual | Near-threshold TMS-induced phosphene perception | Phase-amplitude effect on detection, stronger for high amplitude |
| Other modalities |  |  |  |  |  |  |  |
| Hussain et al., 2019 | EEG | Alpha, Beta | 8-12 Hz, 13-30 Hz | Spontaneous | Motor | Supra-threshold TMS-induced MEP | Optimal phase (leading to increased MEP) reverses between low and high alpha amplitude; No phase-amplitude effect for beta. |
| Madsen et al., 2019 | EEG | Alpha | 7-13 Hz | Spontaneous | Motor | Supra-threshold TMS-induced MEP | No phase-amplitude effect on MEP amplitude |
| Studies not using TMS |  |  |  |  |  |  |  |
| Visual modality |  |  |  |  |  |  |  |
| Alexander et al., 2020 | EEG | Alpha | 5.6-14.4 Hz | Spontaneous | Visual | Detection of near-threshold stimulus through eyes closed | Phase effect on detection for high amplitude |
| Bonnefond & Jensen, 2015 | MEG | Alpha | 9-12 Hz | Stimulus-locked (Retention interval) | Visual | Sternberg working memory task with supra-threshold stimuli | Phase effect on gamma power for high amplitude |
| Busch & VanRullen, 2010 | EEG | Theta | 7 Hz | Spontaneous (amplitude attentionally modulated) | Visual | Detection of spatially attended or unattended near-threshold visual stimulus | Phase effect on detection and GFP for low amplitude |
| Harris et al., 2018 | EEG | Theta, Alpha | 5 Hz, 11-15 Hz | Stimulus-locked (alpha amplitude attentionally modulated) | Visual | Detection of spatially attended or unattended near-threshold visual stimulus | No phase-amplitude effect on detection |
| Kizuk & Mathewson, 2017 | EEG | Alpha | 12 Hz | Entrained | Visual | Detection of spatially attended or unattended metacontrast masked stimulus (75% performance) | Tendency for phase-amplitude effect on detection, stronger for high amplitude |
| Mathewson et al., 2009 | EEG | Alpha | 10 Hz | Stimulus-locked | Visual | Detection of metacontrast masked stimulus | Phase effect on detection for high amplitude |
|  |  |  |  |  |  | (70% performance) |  |
| Milton & Pleydell-Pearce, 2016 | EEG | Alpha | 7.8-12.7 Hz | Spontaneous (amplitude attentionally modulated) | Visual | Simultaneity judgement of spatially attended or unattended stimulus (50% performance) | No phase-amplitude effect on simultaneity judgement |
| Other modalities |  |  |  |  |  |  |  |
| Ai & Ro, 2014 | EEG | Alpha | 8-12 Hz | Spontaneous | Somatosensory | Detection of near-threshold stimulus | Phase effect on detection for high amplitude |
| Herrmann et al., 2016 | MEG | Delta | 2 Hz | Entrained | Auditory | Detection of near-threshold stimulus | Phase effect on detection for high amplitude |
| Ng et al., 2012 | EEG | Theta, Alpha | 2-6 Hz, 8-12 Hz | Entrained | Auditory | Detection of near-threshold stimulus | Theta phase-amplitude effect on detection, stronger for high amplitude; No phase-amplitude effect for alpha. |
| Spitzer et al., 2016 | EEG | Delta, Theta and Alpha | 3 Hz, 6.7 Hz, 11 Hz | Stimulus-locked | Visual, Auditory, and Somatosensory | Two-interval numerosity comparison task (80% performance) | Delta phase-amplitude effect on choice-predictive signals, stronger for high amplitude; No phase-amplitude effect for theta and alpha. |
| Zoefel & Heil, 2013 | EEG | Delta | 0.5 Hz | Entrained and Spontaneous | Auditory | Detection of near-threshold stimulus | No phase-amplitude effect on detection neither for entrained nor for spontaneous oscillations. |

We investigated the causal effect of spontaneous alpha phase-amplitude tradeoffs on cortical excitability and subsequent perceptual performance in the visual modality, and tested the predictions made by the Pulsed Inhibition theory using TMS and EEG in human. Cortical excitability was assessed using phosphene perception induced by single-pulse TMS applied over V1/V2 at perceptual threshold (~50% detection), and its post-pulse evoked activity. Our results validate both predictions, demonstrating that alpha oscillations create periodic inhibitory moments when alpha amplitude is high, leading to lower perceptual performance. Critically, both simulations and ERP analyses confirmed that the obtained results were not a mere analysis confound in which the quality of the phase estimation covaries with the amplitude.

## Materials and Methods

This study is a reappraisal of an early study from Dugué et al. (2011a), which focused on the causal link between the phase of ongoing alpha oscillations, cortical excitability, and visual perception. Here, we instead focused on the combined role of the phase and the amplitude of spontaneous alpha oscillations on the causal relation between cortical excitability and visual perception.

### Participants

As in the original study, the data from nine participants (8 male, 20-35 years-old) were analyzed (see Dugué et al. 2011a, for inclusion/exclusion criteria). All participants fulfilled the standard inclusion criteria for TMS experiment (Rossi et al., 2009), gave their written informed consent, and were compensated for their participation. Human participants were recruited in Toulouse, France. The study was approved by the local French ethics committee “Sud-Ouest et Outre-Mer I” (IRB #2009-A01087-50) and followed the Declaration of Helsinki.

### TMS apparatus and EEG recording

Participants seated in a dark room, 57 cm from a computer screen (36.5° × 27° of visual angle). Their head was maintained using a chinrest and a headrest. A 70-mm figure-of-eight coil was placed over the right occipital pole (V1/V2; ~1 cm above the inion and ~2 cm away from the midline). The handle of the coil was oriented vertically, with the handle of the coil positioned dorsally to the coil itself, resulting in a ventral to dorsal electric current in the brain tissue. Biphasic TMS pulses were applied with a Magstim Rapid^2^ stimulator of 3.5 tesla (Magstim, Spring Garden Whitland, Great Britain). EEG was acquired simultaneously with a 64-channels Active Two Biosemi system (https://www.biosemi.com/index.htm, Amsterdam, The Netherlands), with DC recording at a sampling rate of 1024 Hz. Additional electrodes, CMS (Common Mode Sense) and DRL (Driven Right Leg), were placed 2 cm under the eyes of the participants and were used as reference and ground, respectively, to minimize TMS-induced EEG artifact. Finally, horizontal, and vertical electro-oculograms were recorded using three additional electrodes placed around the eyes.

### Experimental procedure

#### Phosphene screening and titration

Participants were selected based on their ability to perceive TMS-induced phosphenes in the left visual field. A train of 7 pulses at 20 Hz and 70% of the TMS machine output intensity, i.e., suprathreshold, was applied over the right occipital pole (i.e., V1/V2; Dugué et al., 2011a; see also Dugué et al., 2016; 2019; Lin et al., 2021) while participants kept fixating at a central fixation. 24% of the participants did not perceive any phosphene (4 out of 17) and were thus excluded from the main experiment. For each remaining participant, an individual phosphene perception threshold was determined by applying single pulses of TMS at varying intensities and asking them to report (with their dominant hand) whether they perceived a phosphene or not (left or right arrow on the computer keyboard, respectively).

#### Experimental session

Participants performed four blocks of 200 trials each, composed of 90% of test trials and 10% of catch trials (randomly interleaved). In the test trials, single-pulse TMS was applied at the perception threshold – on average across participants, phosphenes were perceived in 45.96% ± 7.68% of trials. In the catch trials, the stimulation intensity was kept the same, but instead of applying single pulses, double pulses (40 ms-interval) were administered to monitor the validity of participants’ responses (91.66% ± 8.81% of phosphene perceived). In other words, participants were presumably not pressing the button randomly; the selected cortical location did in fact lead to phosphene perception when stimulated. Adding a second pulse of TMS with a short delay between the two pulses has a cumulative effect on neural activity, leading to suprathreshold stimulation (Ray et al., 1998; Gerwig et al., 2005; Kammer and Baumann, 2010). The longer the delay between the two pulses, the most likely such cumulative effect disappears, which in the present case would lead for the participants to perceiving two pulses. Debriefing with each participant confirmed that this was never the case. Throughout the experiment, participants kept fixating a central dot and pressed a button to start the trial (**Figure 1**). The delay between the button-press and the subsequent TMS pulse varied randomly between 1500 ms and 2500 ms. After a 600 ms-delay following the pulse, a response screen was displayed instructing the participants to indicate whether they perceived a phosphene or not with the left or right arrow, respectively. The percentage of phosphene perceived was monitored every 20 trials. When above 75% or below 25% of phosphene perceived in the test trials, the experimenter repeated the threshold procedure to select an intensity so as to maintain a threshold around 50% of phosphene perceived.

**Figure 1:**
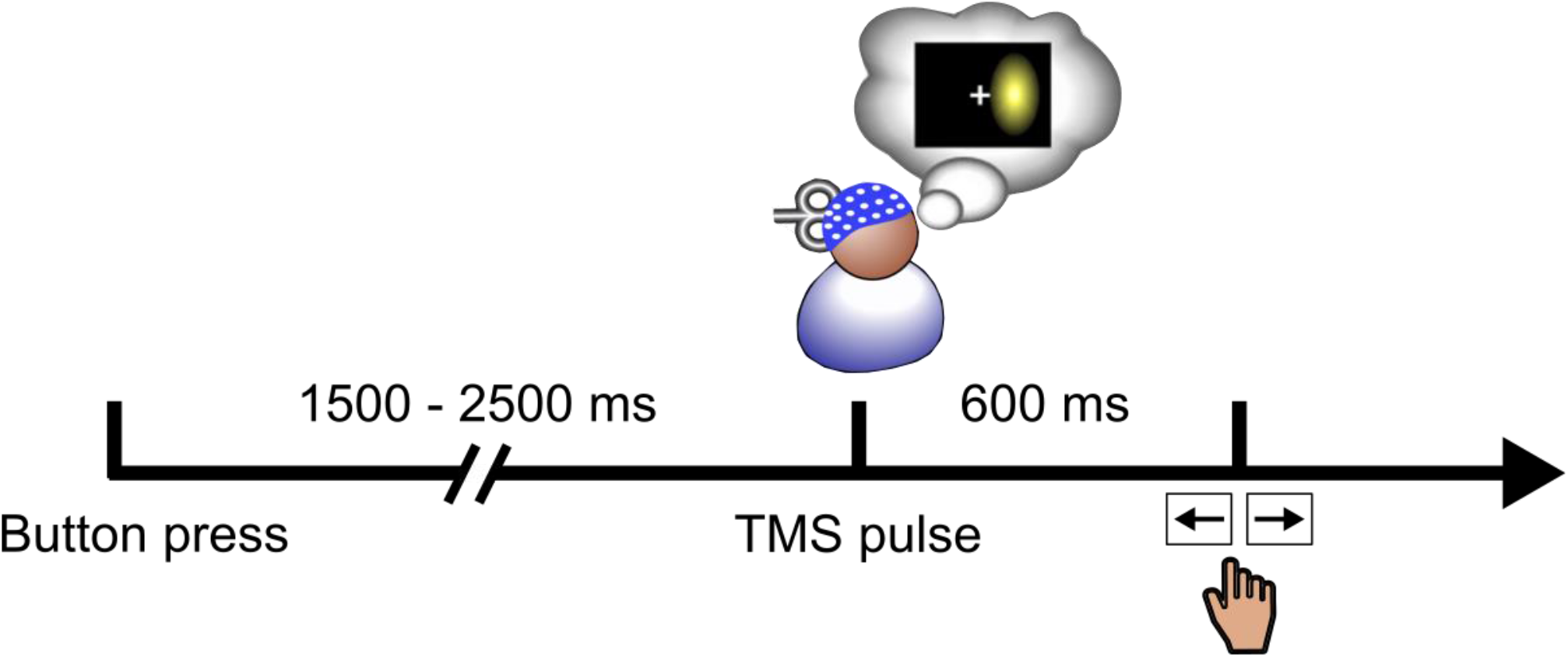
Experimental Paradigm. Participants self-initiated the trial by pressing a button. After a random delay between 1500 and 2500 ms, a single-(90% of trials) or a double-(10% of trials) pulse of TMS was applied over V1/V2. After a 600 ms-delay, participants indicated whether they perceived the phosphene or not with the left or right arrow, respectively.

### EEG Analyses

EEG analyses were performed with EEGLAB 13.6.5 (Swartz Center for Computational Neuroscience, UC San Diego, California; Delorme & Makeig, 2004) and custom software written in MATLAB R2014b (The MathWorks, Natick, MA).

#### EEG preprocessing

EEG data for test trials and channel localizations were imported into EEGLAB. EEG data were downsampled to 512 Hz, re-referenced to average reference, and epoched from −1500 ms to 1000 ms around the single-pulse of TMS. To minimize the artifact induced by the pulse, EEG data from −1 ms to 150 ms around the pulse was erased and replaced with a linear interpolation of the window boundaries. Epochs were finally manually inspected and rejected if artifacts (e.g., blinks) were detected. No electrode was rejected from the analysis.

#### Time-frequency decomposition

A time-frequency transform (morlet wavelets) was computed on single trials with the timefreq function from EEGLAB. The “cycles” parameter was set to [1, 15], the length of the filter increasing logarithmically from 1 to 15 cycles. The “freqs” parameter was set to [2, 100], producing frequencies that increase from 2 to 100 Hz.

#### Trials sorting by bin of amplitude

We then sorted the trials according to the amplitude of pre-pulse, spontaneous, alpha oscillations. The amplitude at each time-frequency point and for each trial, participant, and electrode was thus calculated. It corresponds to the absolute of the complex vector obtained from the time-frequency decomposition. For each trial and participant, we then averaged the amplitude across all electrodes – we had no a priori hypothesis regarding the electrode location of the effect – and time-frequency points selected based on the previous publication (Dugué et al., 2011a), i.e., time-frequency points at which a significant effect of the pre-pulse phase on phosphene perception was observed (from −400 ms to −50 ms before the pulse to avoid post-pulse contamination on pre-pulse activity, and from 7 Hz to 17 Hz). Trials were then sorted in three equally sized bins of low-, medium-, and high-amplitude trials. The percentage of phosphene perceived was computed for each bin of amplitude. Kruskal-Wallis test was used to test for significant difference in phosphene perception across bins. The next analyses were performed on the two extreme bins, i.e., low- and high-amplitude, exclusively, to clearly separate the low- and high-alpha amplitude trials (with no possibility of overlap) using the least number of bins necessary to maximize the number of trials per condition. There were on average across the 9 participants 108 ± 17.92 trials for the perceived- and 115.56 ± 15.86 trials for the unperceived-phosphene condition for the low-alpha amplitude trials, and 98.56 ± 30 trials for the perceived- and 126.11 ± 29.73 trials for the unperceived-phosphene condition for the high-alpha amplitude trials.

#### Phase-opposition

We calculated the phase-locking values (i.e., amount of phase concentration across trials), separately for low- and high-alpha amplitude trials, and perceived- and unperceived-phosphene trials. The phase-locking value was obtained by, first, dividing the complex vectors obtained from the time-frequency decomposition by their length (i.e., instantaneous amplitude) thus normalizing for amplitude and keeping only the instantaneous phase (i.e., angle of the vectors). Second, the mean across trials of the normalized vectors was computed. The length of the average vector is now a measure of phase distribution, i.e., phase-locking across trials. For each amplitude condition, we subsampled the number of trials in the phosphene condition with the most trials to match the phosphene condition with the least trials. This subsampling procedure was repeated 100 times with a different subset of selected trials, and then we averaged the iterations. For each participant, phase-locking values were then summed across perceived- and unperceived-phosphene trials to obtain phase-opposition sums (POS). POS were then averaged across all electrodes, separately for low- and high-alpha amplitude trials.

This measure of phase opposition is designed to give a low value when summing over two conditions both with uniform (random) phase distributions. On the other hand, POS is high when summing over two conditions both with strong phase-locked distributions across trials, as would happen when two conditions are associated with opposite phases. Since POS is computed on spontaneous (pre-pulse) activity, i.e., the phase distribution across all trials can be assumed to be uniform (see below for further statistics), if the phase is locked across trials for one specific condition (half of the overall trials) then the phase of the other condition (other half of the overall trials) would logically be locked in the opposite direction, leading to a high POS value.

This average was then compared with a surrogate distribution obtained with a permutation procedure (Dugué et al., 2015, 2011a; VanRullen, 2016b) consisting in shuffling the perceived- and unperceived-phosphene labels (5,000 repetitions) and recalculating phase-locking values (including subsampling) to obtain a surrogate phase-locking distribution under the null hypothesis that both perceived- and unperceived-phosphene trials have a uniform phase distribution, and further summed across the two surrogate distributions to compute a surrogate POS, characterized by a given mean and standard error of the mean (separate procedure for low- and high-alpha amplitude). Z-scores were computed by comparing the experimentally obtained POS to the mean and standard error of the mean of the surrogate POS.

Z-scores = (POS – surrogate POS mean) / surrogate POS standard error of the mean

FDR correction for multiple comparisons was further applied to p-values. A topographical analysis on the Z-scores revealed two regions of interest (ROI) involved in the phase-opposition effect (occipital and frontal). We repeated the previous analyses for these regions.

Specific time-frequency points were then selected for further analyses. We selected the time points separately for occipital and frontal electrodes/ROIs for which Dugué et al. (2011a) observed the maximal phase effect between perceived- and unperceived-phosphene conditions at electrodes PO3 (in occipital ROI: −77 ms) and AFz (in frontal ROI: −40 ms), respectively. We selected the 10.7 Hz frequency for both occipital and frontal electrodes/ROIs based on the FFT performed on pre-pulse ERPs (see below); this frequency is identical to the frequency obtained when performing FFTs on pre-pulse EEG time series (see next section). Note that the frequency resolution differs between the wavelet decomposition and the FFT. Thus, for all wavelet decomposition related analyses, we used the closest frequency (10.7 Hz) to the peak observed in the FFT analyses (10.24 Hz).

We further ensured, at these selected time-frequency points, that the pre-pulse phase-opposition effect was not due to a contamination by the wavelet decomposition from the post-pulse activity, separately for electrode PO3 and AFz. We thus tested whether the phase was uniformly distributed for both low- and high-alpha amplitude trials. For each trial, phases were extracted and averaged across participants, separately for perceived- and unperceived-phosphene conditions, and the uniformity of the distribution across all trials was tested with a Rayleigh test from the Circular Statistics Toolbox (P. Berens, CircStat: A Matlab Toolbox for Circular Statistics, Journal of Statistical Software, Volume 31, Issue 10, 2009 http://www.jstatsoft.org/v31/i10). For both amplitude conditions, and for both electrode PO3 (low-alpha amplitude trials: p-value = 0.081657, Kappa = 0.1883; high-alpha amplitude trials: p-value = 0.22314, Kappa = 0.13517) and electrode AFz (low-alpha amplitude trials, p-value = 0.65929, Kappa = 0.076575; high-alpha amplitude trials, p-value = 0.19943, Kappa = 0.14016), the tests did not reveal a significant effect suggesting that the phase was uniformly distributed across trials.

Finally, post-hoc one-tailed t-tests were performed to investigate whether POS, computed for each participant at the selected time-frequency points and averaged across electrodes for the occipital and the frontal ROI separately, differed significantly between low- and high-alpha amplitude trials. This analysis was similarly performed for several versions of alpha amplitude binning (i.e., 2, 3, 4 and 5). On average across the 9 participants, phosphene-conditions, and alpha-amplitude conditions, there were 167.94 ± 36.01 trials per bin in the 2-bins version, 111.96 ± 24.29 trials in the 3-bins version, 83.97 ± 18.41 trials in the 4-bins version, and 67.18 ± 15.39 trials in the 5-bins version.

#### Fast Fourier Transform (FFT) on pre-pulse EEG time series

To confirm further that the high-as well as the low-alpha amplitude conditions both contained alpha oscillations, EEG time series from −600 ms to −1 ms were analyzed with an FFT (500 points zero padding), independently for the occipital and frontal ROI, for each alpha-amplitude condition, trial and participant. The resulting amplitude spectra were then averaged across trials and participants and plotted from 2 Hz to 40 Hz. One-tailed t-tests were used to compare individual 10.24 Hz peaks to their corresponding 1/f aperiodic component. The 10.24 Hz peaks were further compared between low- and high-alpha amplitude conditions with one-tailed t-tests. To ensure that the significant difference between the two conditions did not come from a difference in their aperiodic 1/f component, we also fitted the amplitude spectra to the 1/f component for each participant and compared the 1/f component at 10.24 Hz between low- and high-alpha amplitude conditions with two-tailed t-tests.

#### Simulations

The phase-opposition analysis between low- and high-alpha amplitude trials could be due to an analysis confound, i.e., with decreasing amplitude, the robustness of the phase estimation decreases. Hence, a control procedure ensured that the phase estimation was not impacted by amplitude covariation, especially relevant when interpreting phase-opposition in low-alpha amplitude trials. A time-frequency decomposition and phase-opposition analysis, identical to the one described above (**Figure 2**), was performed on a simulated dataset. Four electrophysiological datasets were simulated with similar properties as those observed in our empirical data: one for each experimental condition (300 trials each), i.e., perceived- and unperceived-phosphene for low- and high-alpha amplitude conditions. Specifically, each trial was created as a sum of sine waves from 2 to 40 Hz. The amplitude of the simulated signal was determined based on the empirical data. For frequencies from 2-7 Hz and 13-40 Hz, an amplitude of 100 au (arbitrary unit) was chosen. For frequencies from 8-12 Hz, amplitudes varied depending on the simulated condition. For the low-alpha amplitude condition, we selected amplitudes of 280 au and 320 au for perceived- and unperceived-phosphene condition, respectively. For the high-alpha amplitude condition, we selected amplitudes of 580 au and 620 au for perceived- and unperceived-phosphene conditions, respectively. A random phase between 0 and 2π was selected for frequencies from 2-7 Hz and 13-40 Hz. For frequencies from 8-12 Hz, the phase varied depending on the simulated condition, i.e., [0 π] for perceived-phosphene for both low- and high-alpha amplitude conditions, and [π 2π] for unperceived-phosphene for both low- and high-alpha amplitude conditions. This phase distribution was applied to 80% of trials. In the other 20% of trials, a random phase between 0 and 2π was selected. Finally, white noise (VanRullen, 2016b) was added to all simulated datasets (amplitude: 4000 au) to match the empirically observed averaged z-score of phase-opposition (**Figure 2**).

**Figure 2.**
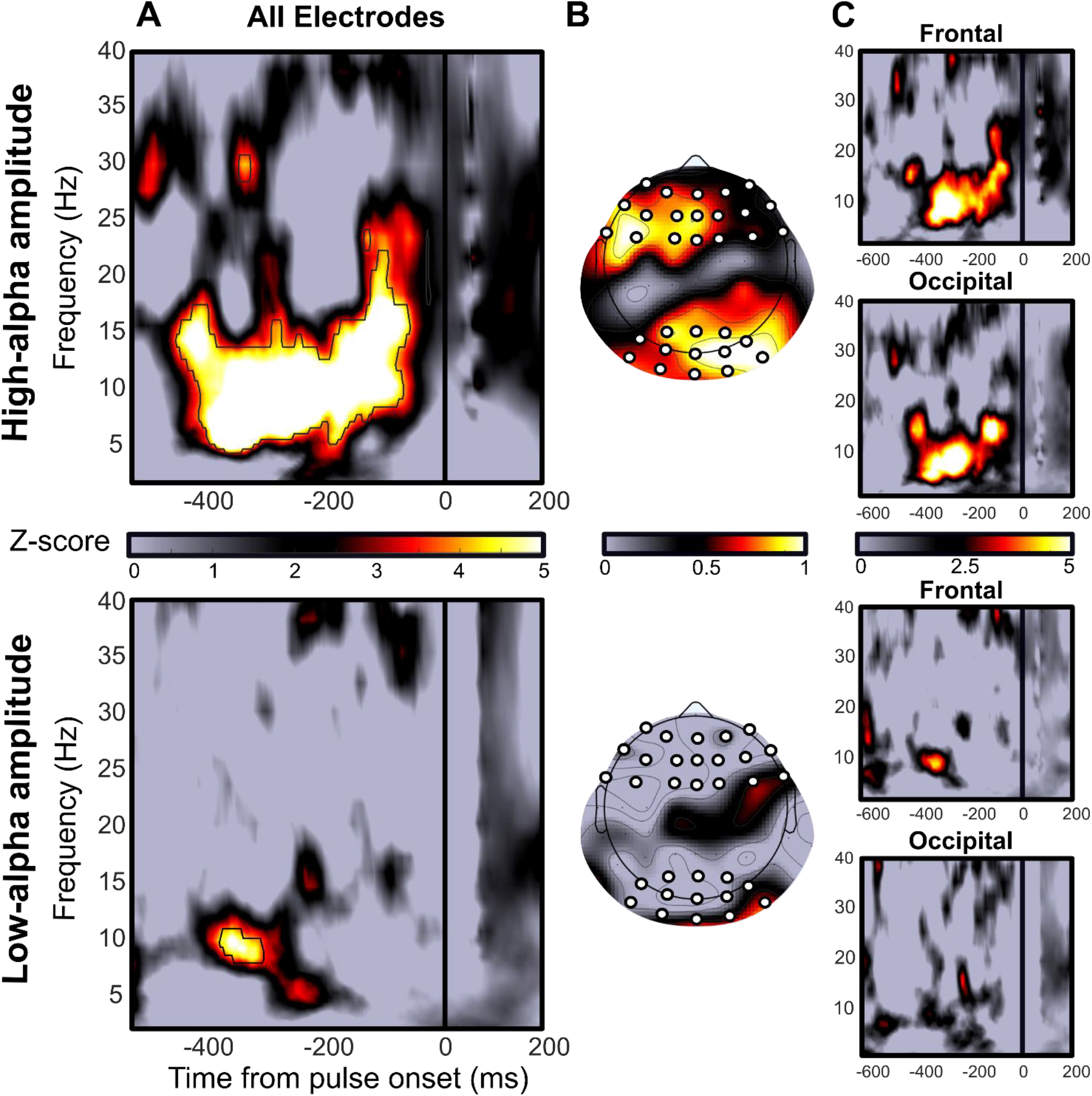
The phase of spontaneous alpha oscillations predicts phosphene perception mainly for high-alpha amplitude. This figure is supported by Extended Data Figures 2-1, 2-2 and 2-3. Upper panel, phase-opposition computed respectively on high-alpha amplitude trials. Lower panel, low-alpha amplitude trials. **A.** Z-scores map of phase-opposition between perceived- and unperceived-phosphene conditions averaged across 9 participants and all 64 electrodes. Colormap, Z-scores. Black outline, significant phase-opposition FDR corrected for multiple comparisons (FDR = 0.01, corresponding to p-values threshold of 2.08×10^−5^ for low-alpha amplitude condition, and 1.96×10^−5^ for high-alpha amplitude condition). There is a significant phase-opposition from −400 ms to −50 ms before the pulse, between 5 and 18 Hz when alpha amplitude is high. The effect is less extended across time and frequencies when alpha amplitude is low. **B.** Z-scores topographies averaged across the time-frequency window identified in panel A for high-alpha amplitude. The effect is maximal in a frontal and an occipital region-of-interest (ROI) when alpha amplitude is high. The topography is less clear when alpha amplitude is low. White dots, electrodes of interest within each ROI. **C.** Z-scores maps of phase-opposition computed separately for the frontal ROI (upper panel) and the occipital ROI (lower panel).

**Extended Data Figure 2-1.**
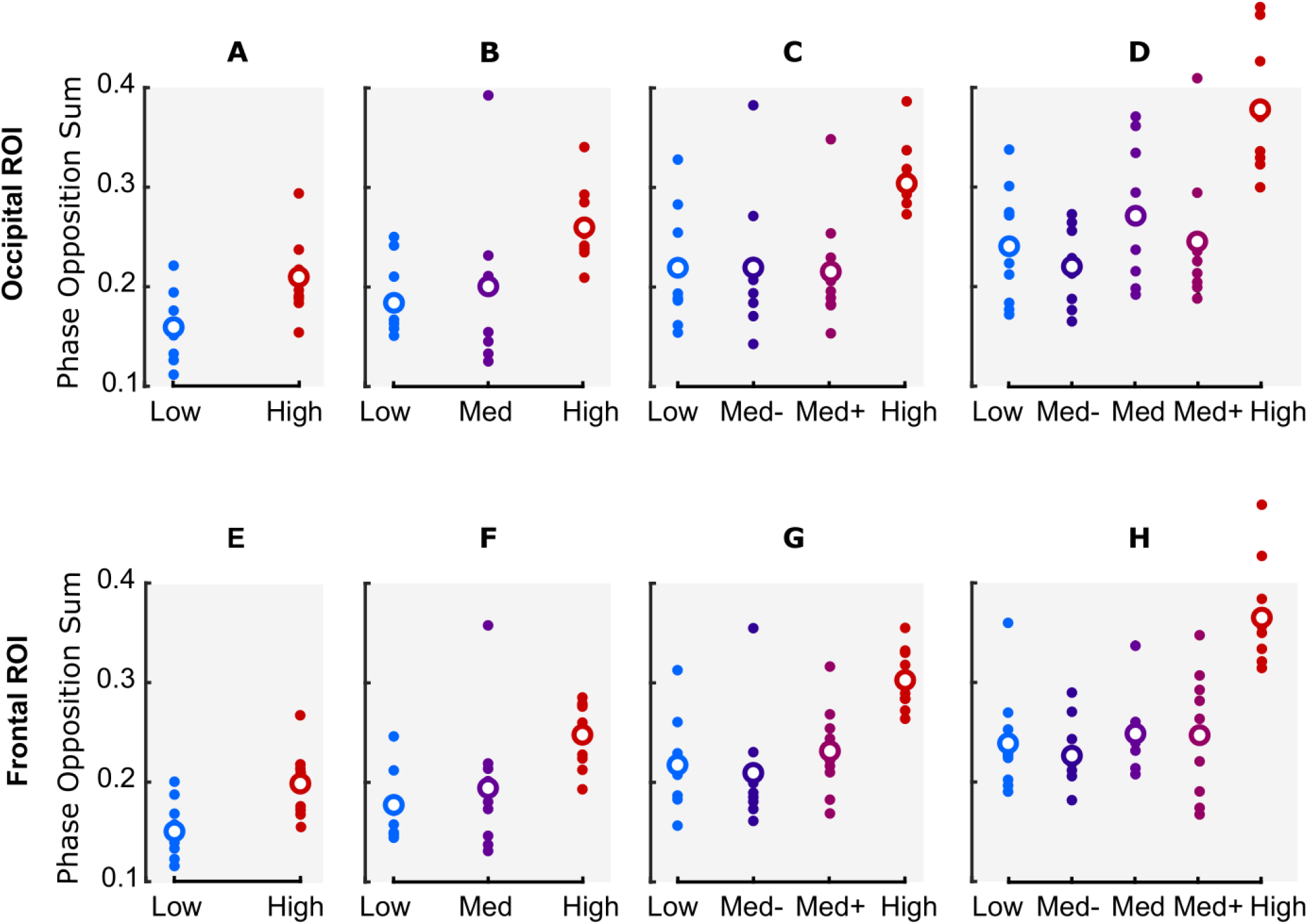
The phase effect on phosphene perception is higher for high-compared to low-alpha amplitude trials. Phase-opposition sum computed for several binning versions of alpha amplitude trials, at 10.7 Hz and **A-D** at −77 ms pre-pulse, and averaged across electrodes within the occipital ROI, **E-H** at −40 ms pre-pulse, and averaged across the electrodes within the frontal ROI. Dots, POS for individual participants. Circles, POS averaged across the 9 participants. All following analyses were performed on the 3-bins condition (**B** and **F**), i.e., trials were binned in low-, medium- (med) and high-alpha amplitude trials, discarding the medium-alpha amplitude bin.

**Extended Data Figure 2-2.**
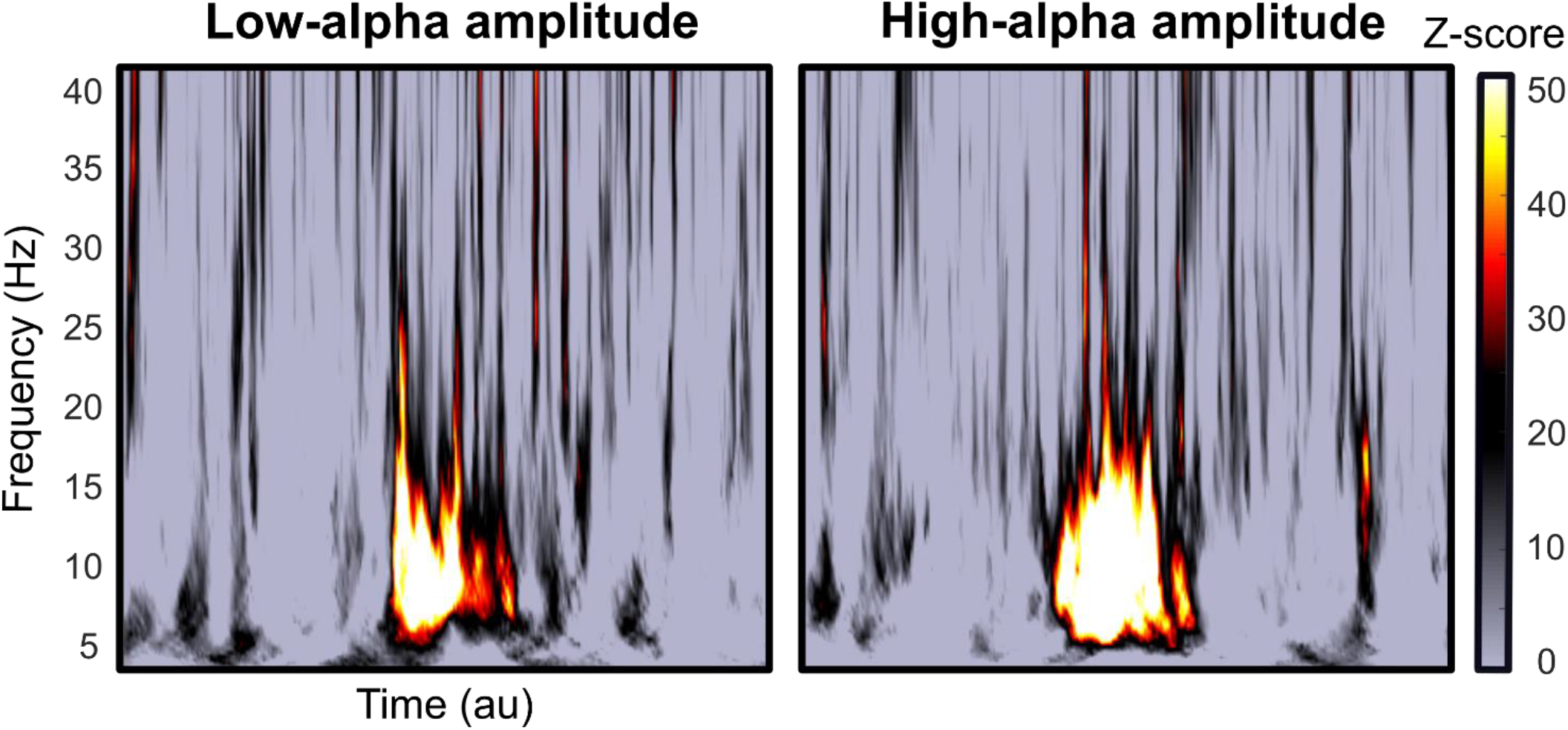
Both low- and high-alpha amplitude oscillations simulated datasets show a similar phase effect on perception. Left panel, phase-opposition computed on simulated low-alpha amplitude trials. Right panel, simulated high-alpha amplitude trials. Z-scores maps of phase-opposition between perceived- and unperceived-phosphene conditions. Colormap, Z-scores. Au, Arbitrary Unit. Between low- and high-alpha amplitude simulated trials, there is a comparable phase-opposition between perceived- and unperceived-phosphene conditions, from 5 to 18 Hz.

**Extended Data Figure 2-3.**
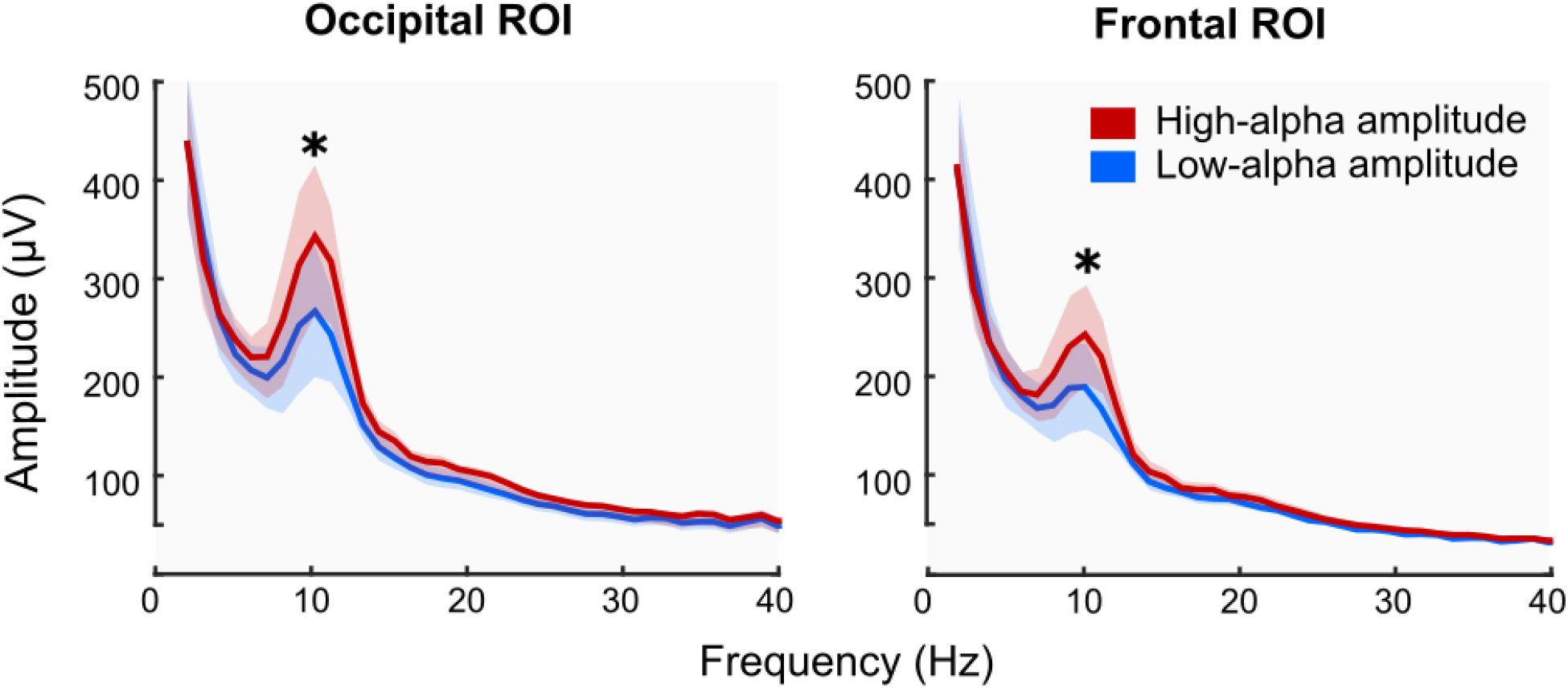
Pre-pulse oscillatory activity in the alpha frequency band in both low- and high-alpha amplitude conditions. Amplitude spectra computed on the EEG time-series from −600 ms to −1 ms relative to pulse onset, for the occipital ROI (left panel) and the frontal (right panel) ROI. Red color, high-alpha amplitude condition. Blue color, low-alpha amplitude condition. Colored solid lines, amplitude spectra averaged across the 9 participants between 2 and 40 Hz. Colored shaded areas, standard error of the mean. *, significant difference at 10.24 Hz between low- and high-alpha amplitude conditions.

#### Event Related Potentials (ERPs)

Previously preprocessed EEG data were further cleaned from the power line noise by applying a notch filter at 50 Hz (band-stop at 47-53 Hz) before epoching. ERPs, centered on the pulse onset, were computed as the average of trials for each alpha-amplitude condition, phosphene-perception condition, and participant. The difference of ERPs between perceived- and unperceived-phosphene conditions was computed, separately for low- and high-alpha amplitude conditions. The ERP differences were then compared against zero with repeated measures one-tailed t-tests (from 350 ms to 800 ms). Correction for multiple comparisons was applied following a cluster procedure. For each participant, surrogate ERPs were obtained by shuffling the perceived and unperceived phosphene labels (500 repetitions). T-tests were recomputed similarly as before, and the number of consecutive significant time points (surrogate cluster size) for each repetition was stored. The p-value for each empirical cluster was then computed as the proportion of surrogate clusters that were larger than the empirical cluster. Using an α level of 0.05, we considered an empirical cluster significant if its size was larger than at least 95% of the surrogate clusters.

#### FFT on pre-pulse ERPs

The use of TMS to induce phosphene perception rather than an external stimulation allows for direct access to the instantaneous state of the spontaneous brain oscillations. In other words, one can directly compute the pre-pulse ERP differences (time before pulse onset) between perceived- and unperceived-phosphene conditions, to assess spontaneous oscillatory activity. In this case, when comparing ERP differences between perceived- and unperceived-phosphene conditions, separately for low- and high-alpha (pre-sorted) amplitude trials, there is no analysis confound coming from a less accurate phase estimation when alpha-amplitude is low. ERP differences from −400 ms to 0 ms were then analyzed with an FFT (500 points zero padding), independently for electrodes PO3 and AFz, for each alpha-amplitude condition and for each participant. The resulting amplitude spectra were then averaged across participants and plotted from 2 Hz to 40 Hz (a peak at 10.24 Hz was observed for both electrodes and for each alpha-amplitude condition). We then performed the following steps to test for a difference between high- and low-alpha amplitude trials in the resulting amplitude spectra: 1) difference of the averaged amplitude spectra between high- and low-alpha amplitude trials, 2) fit of this difference to the 1/f component, 3) removing of the 1/f component, 4) gaussian fit of the resulting amplitude spectra to extract the frequency window showing a difference between high- and low-alpha amplitude trials, 5) statistical comparison of the amplitude spectra (uncorrected for 1/f) between high- and low-alpha amplitude trials conditions with a one-tailed t-test. To ensure that the obtained significant difference between low- and high-alpha amplitude conditions was not due to a difference in their 1/f aperiodic components, we fitted the amplitude spectra to their 1/f component and computed the area under the curve, for each participant, separately for low- and high-alpha amplitude conditions, for both electrodes PO3 and AFz. The area under the curve of the low-alpha amplitude condition was compared to the one of the high-alpha amplitude condition with a two-tailed t-test. Finally, an FFT (500 points zero padding) was performed separately for the ERP of perceived- and unperceived-phosphene conditions on each participant. We then computed phase-locking values across participants to assess the inter-individual variability at 10.24 Hz for low- and high-alpha amplitude trials separately (frequency at which a peak of amplitude was observed in the amplitude spectra of the ERP difference between phosphene perceived and unperceived conditions). The sum of phase-locking values across participants of perceived- and unperceived-phosphene conditions was computed, for low- and high-alpha amplitude trials separately, and evaluated statistically with a permutation procedure. P-values were estimated by comparing these phase-opposition sums to the mean and standard error of the mean of the surrogate distribution obtained by shuffling the perceived- and unperceived-phosphene labels (repeated 500 times), for low- and high-alpha amplitude trials separately, before replicating the previous analysis on the surrogate ERPs.

#### FFT on post-pulse ERPs

As mentioned above, the advantage of the present TMS procedure is that it allows for direct access to the instantaneous state of the spontaneous brain oscillations. In other words, pre-pulse spontaneous activity is readily observable on the ERP. To ensure that post-pulse ERPs were not contaminated by spontaneous alpha oscillations (i.e., that the TMS pulse here reset alpha oscillations), an FFT (500 points zero padding) was performed on the ERPs of the perceived- and unperceived-phosphene conditions, from 400 ms to 800 ms, for electrodes PO3 and AFz separately, and for each alpha-amplitude condition and participant. One-tailed t-tests against the aperiodic 1/f activity were used to test the significance of the 10.24 Hz peak.

#### Perceptual performance as a function of pre-pulse phase

Low- and high-alpha amplitude trials were sorted in 9 phase bins at the selected time-frequency points (see **Phase-opposition** section), separately for the occipital ROI, as well as the specific electrode PO3 (−77 ms, 10.7 Hz), and the frontal ROI, as well as the specific electrode AFz (−40 ms, 10.7 Hz). The percentage of perceived-phosphene was computed for each phase bin, alpha-amplitude condition, electrode, and participant, and further averaged across participants and electrodes. The values were finally normalized by dividing the percentage of perceived-phosphene averaged across phase bins, separately for each alpha-amplitude condition. A two-way repeated-measures ANOVA was performed to test for the main effect of phase bin. The percentage of variance explained was computed as the difference between the optimal phase (maximum percentage of phosphene perceived) and the opposite one.

#### ERP amplitude as a function of pre-pulse phase

Bin sorting was applied as described in the previous section. Then, the ERP difference between perceived- and unperceived-phosphene conditions for each phase bin and alpha-amplitude condition was computed. The maximum perceived-unperceived ERP difference was selected in the time window in which an ERP difference between low- and high-alpha amplitude was detected, according to repeated measures two-tailed t-test for each time point from 350 ms to 800 ms (PO3: from 482 ms to 513 ms; AFz: from 605 ms to 728 ms). A two-way repeated-measures ANOVA was performed to test for a main effect of phase bin and interaction between phase bin and alpha-amplitude (results regarding the main effect of alpha-amplitude were not interpreted). Specifically, we tested the hypotheses that (1) the ERP difference between perceived- and unperceived-phosphene conditions depended on the phase of spontaneous alpha oscillation, and (2) that this phase effect is stronger for high-alpha amplitude trials. Finally, we calculated the maximum ERP amplitude of the perceived- and unperceived-phosphene conditions separately, independently for phase bin centered on −π/4 and π/2, corresponding respectively to the maximum and the minimum ERP difference, and low- and high-alpha amplitude trials. For each alpha-amplitude condition, we fitted the data to a linear mixed-effect model with phase bins and phosphene conditions as fixed effects, and participants as random effects. Post-hoc analyses were done with one-tailed t-tests.

## Results

Single-pulse TMS was applied over the right occipital cortex (V1/V2) in 9 healthy participants, at threshold intensity (45.96% ± 7.68% of phosphene perceived across participants) while simultaneously recording EEG. Previous analysis of this dataset (Dugué et al., 2011a) revealed that the phase of spontaneous alpha oscillations in the time-frequency window from −400 ms to −50 ms pre-pulse, and from 7 Hz to 17 Hz, predicts the perceptual outcome. This phase effect explained ~15% of the variability in phosphene perception. Here, trials were split according to low-, medium-, and high-amplitude of the pre-pulse spontaneous alpha oscillations (within the same time frequency-window as in Dugué et al., 2011a). We tested the two predictions made by the Pulsed Inhibition theory: (1) high alpha amplitude induces periodic inhibitory moments leading to poor perceptual performance; and (2) low alpha amplitude is less susceptible to phasic inhibition, and lead to overall higher perceptual performance.

We first analyzed phosphene detection for each alpha-amplitude condition. We observed that phosphene detection rate depends on alpha amplitude with the highest detection being in low-(48.22% ± 4.88%) then medium-(45.94% ± 7.92%), and high-(43.75% ± 11.57%) alpha amplitude trials (Kruskal-Wallis: p-value = 0.0553, Cohen’s d (effect size) = 0.88). Note that the earlier study (Dugué et al., 2011a) was not optimized to test the predictions made by the Pulsed Inhibition theory. However, the present results argue in its favor, with higher phosphene perception when alpha amplitude is low (Romei et al., 2008). In the next analyses, we discarded the medium-alpha amplitude trials to concentrate on the low- and high-alpha amplitude conditions. This allowed us to clearly separate the two types of amplitude trials, while maximizing the number of trials per condition (see also **Extended Data Figure 2-1** for further assessment of such amplitude binning procedure).

To investigate the potential joint effect of the amplitude and the phase of spontaneous alpha oscillations on phosphene perception, we calculated phase-opposition sum (POS; see **Materials and Methods** section), separately for low- and high-alpha amplitude conditions. Specifically, this analysis assesses whether phosphene perception is modulated by significantly different phases of the alpha cycle by computing the sum of phase-locking values (i.e., the amount of phase concentration across trials) over the perceived- and unperceived-phosphene conditions (Dugué et al., 2011a, 2011b, 2015, VanRullen, 2016b). Spontaneous activity is characterized by a uniform phase distribution across all trials. Thus, if the phase is locked across trials for the perceived-phosphene condition (approximately half of the overall trials) then the phase of the unperceived-phosphene condition (other half of the overall trials) will logically be locked in the opposite direction, leading to a strong POS. Conversely, a weak POS value can only be obtained if both perceived and unperceived-phosphene conditions have near-random phase distributions, i.e., if phase does not affect phosphene perception. For high-alpha amplitude (**Figure 2**, top raw), this analysis revealed a strong phase-opposition between perceived- and unperceived-phosphene conditions across all participants and electrodes, from −400 ms to −50 ms pre-pulse, and in the frequency range from 5 Hz to 18 Hz (**Figure 2A**). This effect remained significant after FDR correction for multiple comparisons (FDR = 0.01, corresponding to a Z-score threshold of 4.27, a p-value threshold of 1.96 × 10^−5^, and a Cohen’s d threshold approaching infinity). The corresponding topography revealed that the phase-opposition effect was maximal over occipital and frontal electrodes (**Figure 2B,C**). The analysis was replicated on low-alpha amplitude trials (**Figure 2**, bottom raw). The overall strength of the effect was less important, i.e., effect less extended across time and frequency, for low-alpha amplitude trials (remained significant after FDR correction, FDR = 0.01, corresponding to a Z-score threshold of 4.26, a p-value threshold of 2.08 × 10^−5^, and a Cohen’s d approaching infinity), and showed a less informative topography of the effect.

A post-hoc analysis revealed that POS values, averaged across electrodes, separately for the occipital and frontal ROIs at the respective selected time-frequency points (see **Materials and Methods**), were significantly higher for high-compared to low-alpha amplitude trials at (10.7 Hz, −77 ms) for the occipital ROI (one-tailed t-test: p-value = 0.0016, Cohen’s d = 1.9645, CI = [0. 0.042; Infinity]) (**Extended Data Figure 2-1B**) and at (10.7 Hz, −40 ms) for the frontal ROI (one-tailed t-test: p-value < 0.001, Cohen’s d = 2.1165, CI = [0. 0.0457; Infinity]) (**Extended Data Figure 2-1F**). Together, these results suggest that there is an optimal phase of spontaneous alpha oscillations that predicts phosphene perception. This phase effect is more robust across time and across frequencies and is significantly higher for high-compared to low-alpha amplitude trials.

Interestingly, **Extended Data Figure 2-1** further illustrates the impact of several binning versions of the previous analysis (from 2 to 5 bins). In all versions, there is a difference of POS between low- and high-alpha amplitude trials (one-tailed t-tests for all binning versions show p-values < 0.004 and Cohen’s ds >1.27). However, increasing the number of bins decreases the number of trials in each bin. Consequently, all main analyses were performed on the 3-bins version (excluding the middle bin) to clearly separate low- and high-alpha amplitude trials while maximizing the number of trials per condition.

To ensure that the difference in phase effect observed between low- and high-alpha amplitude trials does not depend on a poor estimation of the phase when alpha amplitude is low, we performed a control analysis based on simulations (**Extended Data Figure 2-2**). The phase-opposition analysis displayed in **Figure 2** was repeated on simulated data generated with the same parameters (amplitude ratio between low- and high-alpha amplitude trials) than those observed in the empirical dataset (see **Materials and Methods** for more details). The simulations show that in both the low- and high-alpha amplitude conditions, POS between perceived- and unperceived-phosphene trials can be observed with similar time-frequency profiles. Thus, the effect observed in **Figure 2** cannot be simply explained by an underpowered phase estimation in low-alpha amplitude trials but indeed reflects a functional neurophysiological brain process.

In addition, we tested that alpha oscillations are actually present in pre-pulse, low-alpha amplitude trials (**Extended Data Figure 2-3**). An FFT on the pre-pulse EEG activity revealed a peak at 10.24 Hz in the occipital ROI for both low-(one-tailed t-test against the 1/f aperiodic activity: p-value =0.0354, Cohen’s d = 0.7371, CI = [11.7481; Infinity]) and high-(p-value = 0.0102, Cohen’s d = 0.9999, CI = [60.0927; Infinity]) alpha amplitude trials, and in the frontal ROI for both low-(p-value = 0.0465, Cohen’s d = 0.6668, CI = [1.6258; Infinity]) and high-(p-value = 0.0162, Cohen’s d = 0.9527, CI = [29.2137; Infinity]) alpha amplitude trials. This analysis confirms that estimating the phase in low-alpha amplitude trials is indeed neurophysiologically relevant. Additionally, the peak at 10.24 Hz was significantly higher for high-compared to low-alpha amplitude trials, for both the occipital (one-tailed t-tests: p-value = 0.0268, Cohen’s d = 0.2176, CI = [6.8434; Infinity]) and the frontal (p-value = 0.0316, Cohen’s d = 0.2485, CI = [4.1668; Infinity]) ROI. This difference was unlikely due to a difference in the 1/f aperiodic activity between low- and high-alpha amplitude conditions, i.e., there was no significant difference in the 1/f component at 10.24 Hz between low- and high-alpha amplitude conditions for the occipital (two-tailed t-tests: p-value = 0.2362; Cohen’s d = 0.1893, CI = [−9.8442; 34.4352]) and the frontal (p-value = 0.0867, Cohen’s d = 0.3361, CI = [−2.5463; 30.6516]) ROI.

To further understand the link between pre-pulse spontaneous alpha oscillatory phase and amplitude, cortical excitability and phosphene perception, and address further a possible confound coming from a less accurate phase estimation when alpha amplitude is low (see **Materials and Methods** section), we analyzed the ERP difference between perceived- and unperceived-phosphene conditions. Critically, the use of TMS to induce phosphene perception allows for direct access to the instantaneous state of the spontaneous brain oscillations. In other words, the pre-pulse ERP differences (time before pulse onset) between perceived- and unperceived-phosphene conditions, allows to assess spontaneous oscillatory activity. Thus, if there is an optimal phase for perception and an opposite, non-optimal one, then for each participant the pre-pulse ERP for perceived-phosphene and for unperceived-phosphene should each oscillate in alpha, and so would the ERP difference. Additionally, if all participants share the same optimal phase, then the pre-pulse ERP difference averaged across participants should oscillate in alpha as well. We analyzed the ERP difference between perceived- and unperceived-phosphene conditions for electrodes PO3 and AFz (selected respectively in the occipital and frontal ROIs based on previous studies; Taylor et al., 2010; Dugué et al., 2011a; **Figure 3**). For both electrodes and for both low- and high-alpha amplitude trials, the ERP difference appeared periodic in the last 400 ms preceding the pulse (note that both low- and high-alpha amplitude conditions show this effect). An FFT applied on the ERP difference of each participant in the pre-pulse period (−400 to 0 ms) showed a peak in amplitude for both electrode PO3 (**Figure 4A;** 10.24 Hz for both low- and high-alpha amplitude) and electrode AFz (**Figure 4B;** 10.24 Hz for low- and 9.22 Hz for high-alpha amplitude). Additionally, the amplitude of the pre-pulse oscillatory difference between perceived- and unperceived-phosphene was significantly higher for high-alpha amplitude compared to low-alpha amplitude trials, for the frequency window from 5.12 Hz to 11.26 Hz for PO3 (one-tailed t-test: p-value = 0.044, Cohen’s d = 0.361, CI = [0.813; Infinity]; see **Materials & Methods** section), and from 7.16 Hz to 10.24 Hz for AFz (p-value = 0.039, Cohen’s d = 0.241, CI = [0.786; Infinity]). This difference was unlikely due to a difference in the 1/f aperiodic activity between low- and high-alpha amplitude conditions as their aperiodic activity did not differ significantly, neither for electrode PO3 (two-tailed t-test: p-value = 0.063, Cohen’s d = 0.3131, CI = [−13.8069; 414.8763]) nor AFz (two-tailed t-test: p-value = 0.806, Cohen’s d = 0.0264, CI = [−76.8006; 95.8066]). To further assess the inter-individual variability, we calculated the sum of phase-locking values across participants at 10.24 Hz for perceived- and unperceived-phosphene conditions, separately for low- and high-alpha amplitude trials, and for PO3 and AFz electrodes. In other words, we ask whether the pre-pulse ERP for each condition oscillates in-phase across all participants, and are in phase-opposition between perceived- and unperceived-phosphene conditions. We found that there is a phase-opposition between perceived- and unperceived-phosphene conditions for the electrode PO3, for high-alpha amplitude trials (permutation statistics: Z-score = 1.9794, p-value = 0.0239, Cohen’s d = 1.756), but not for low-alpha amplitude trials (Z-score = 0.9856, p-value = 0.1622, Cohen’s d = 0.6957), nor for AFz low-(Z-score = 0.2059, p-value = 0.4184, Cohen’s d = 0.1375) and high-(Z-score = 1.169, p-value = 0.1212, Cohen’s d = 0.8462) alpha amplitude. Phosphene perception depends on an optimal phase of alpha oscillation at the occipital electrode PO3, when alpha amplitude is high. Together, these analyses suggest that phosphene perception alternates between optimal and non-optimal phases of the alpha (10.24 Hz) oscillations in the 400 ms window before the pulse, with all participants sharing a similar optimal phase. This phase effect is predominant in the occipital region, and stronger when the alpha amplitude is high.

**Figure 3.**
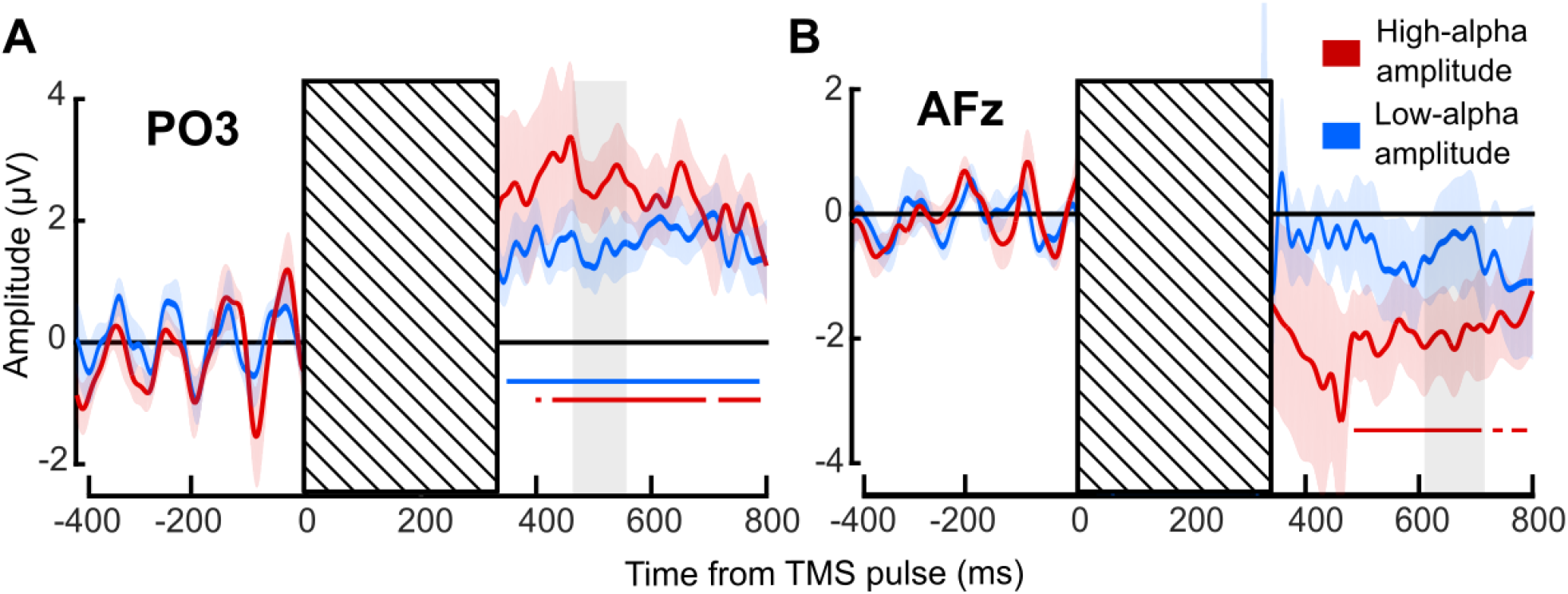
Difference between perceived-phosphene ERP and unperceived-phosphene ERP. This figure is supported by Extended Data Figure 3-1. **A.** ERP difference at electrode PO3 between perceived- and unperceived-phosphene conditions averaged across 9 participants. **B.** ERP difference at electrode AFz. Red, high-alpha amplitude trials. Blue, low-alpha amplitude trials. Colored shaded areas, standard error of the mean. Striped areas, mask the TMS-induced artifact. Colored solid horizontal lines, significant ERP difference against zero (significant cluster for PO3, low- and high-alpha amplitude and AFz, high-alpha amplitude; p<0.001). Gray shaded areas, selected time window of interest.

**Extended Data Figure 3-1.**
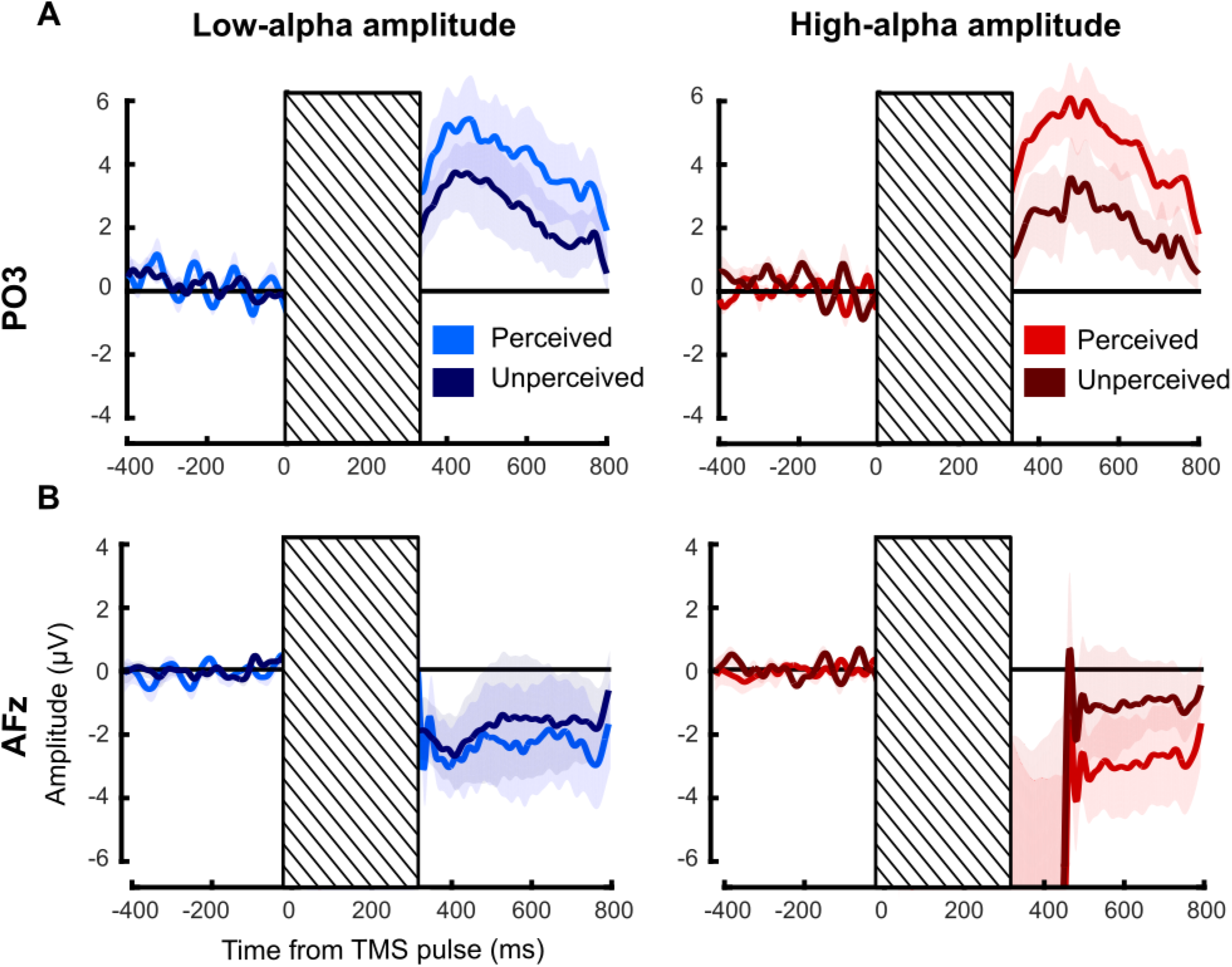
ERPs for perceived- and unperceived-phosphene trials. **A.** ERPs at electrode PO3 for perceived- and unperceived-phosphene trials averaged across the 9 participants. **B.** ERPs at electrode AFz. Red, high-alpha amplitude condition. Blue, low-alpha amplitude condition. Light colors, perceived-phosphene condition. Dark colors, unperceived-phosphene condition. Colored shaded areas, standard error of the mean. Striped area, mask the TMS-induced artifact.

**Figure 4.**
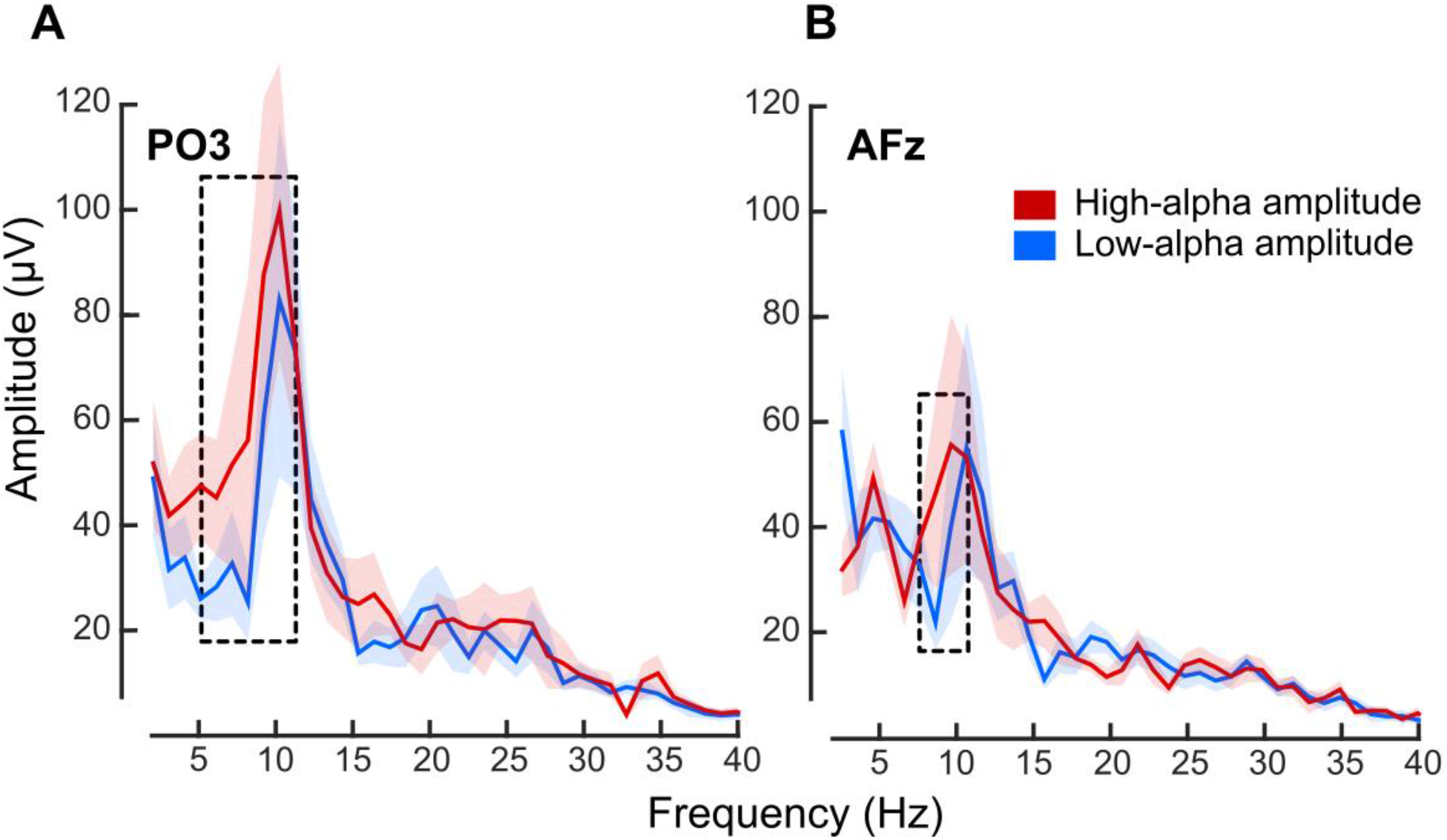
The pre-pulse ERP difference between perceived- and unperceived-phosphene conditions oscillates in alpha. **A.** Frequency spectra computed on the ERP difference between perceived- and unperceived-phosphene conditions, on the pre-pulse period from −400 ms to 0 ms, respectively for electrode PO3, and **B.** electrode AFz. Red color, high-alpha amplitude trials. Blue color, low-alpha amplitude trials. Colored solid lines, frequency spectra averaged across the 9 participants between 2 and 40 Hz. Shaded area, standard error of the mean. Dotted rectangle, significant difference between high- and low-alpha amplitude trials averaged across frequency window from 5.12 to 11.26 Hz for electrode PO3, and from 7.16 to 10.24 Hz for electrode AFz (one-tailed t-tests: p-value = 0.044 for PO3, p-value = 0.039 for AFz). Frequency peak at 10.24 Hz for both low- and high-alpha amplitude for electrode PO3; at 10.24 Hz for low- and 9.22 Hz for high-alpha amplitude for electrode AFz.

Next, for each participant, we sorted low- and high-alpha amplitude trials in 9 bins according to the pre-pulse EEG phase, for the selected time-frequency points (occipital ROI: −77 ms, 10.7 Hz; frontal ROI: −40 ms, 10.7 Hz). Then, the percentage of phosphene perceived was calculated for each bin and averaged across participants (**Figure 5**). A two-way repeated-measures ANOVA revealed a significant effect of the phase in both the occipital (F(1,8) = 2.117, p-value = 0.0467, eta^2^ (effect size) = 20.93, SS (square sum) = 0.345) and frontal (F(1,8) = 3.360, p-value = 0.0028, eta^2^ = 29.58, SS = 0.472) ROIs. There was no main effect of amplitude in either the occipital (F(1,8) = 1.818, p-value = 0.2145, eta^2^ = 18.51, SS = 0.001) or the frontal (F(1,8) = 0.225, p-value = 0.6476, eta^2^ = 2.74, SS < 0.001) ROIs, nor interaction in either the occipital (F(1,8) = 0.249, p-value = 0.9794, SS = 0.048) or the frontal (F(1,8) = 0.462, p-value = 0.8781, SS = 0.076) ROIs. Critically, the results show that the optimal phase for phosphene perception is centered on π/2 while the opposite phase, between −π/2 and −π/4, is non-optimal. Finally, we observe that the percentage of variance explained by the phase is more important for high-(occipital: 16.9% difference between π/2 and −π/2; frontal: 21.2%) compared to low-(occipital: 13.3%; frontal: 17.6%) alpha amplitude of spontaneous oscillations.

**Figure 5.**
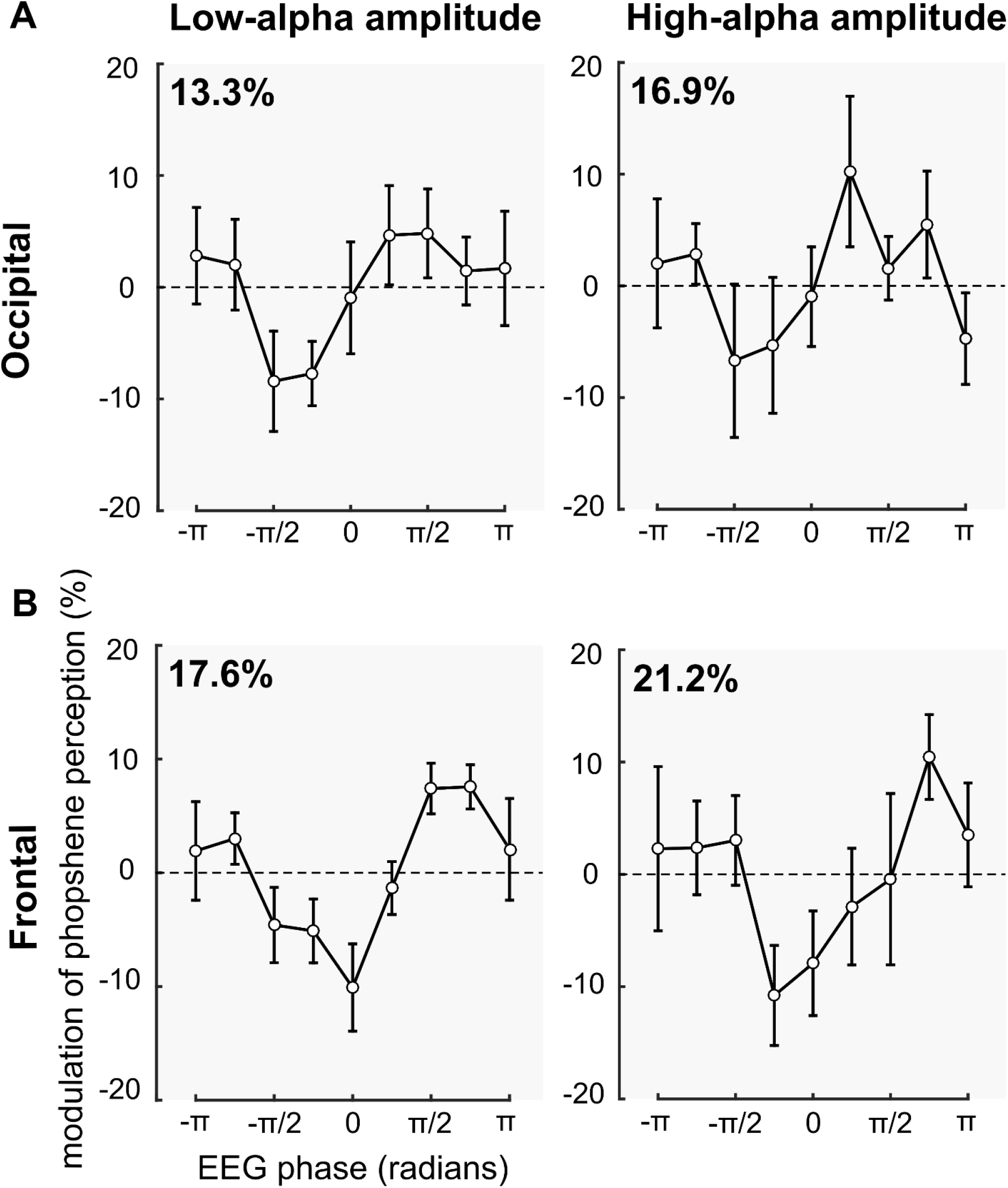
The phase π/2 of the alpha cycle is the optimal phase for phosphene perception. Left panels, phosphene perception computed for 9 phase bins (expressed in radians), normalized according to the average phosphene perception, and averaged across the 9 participants and electrodes of interest, for low-alpha amplitude trials. Right panels, for high-alpha amplitude trials. Error bars, standard error of the mean. **A.** Phosphene perception is plotted according to the instantaneous phase at −77 ms, 10.7 Hz, for the occipital ROI. Phosphene perception oscillates along with the alpha phase (two-way repeated-measures ANOVA: F (1,8) = 2.117, p-value = 0.0467 eta^2^ = 20.93), with an optimal phase for phosphene perception at the phase π/2 of the alpha cycle. The percentage of variance explained by the phase is more important for high-(16.9% difference between π/2 and −π/2) compared to low-(13.3%) alpha amplitude trials. **B.** Phosphene perception is plotted according to the instantaneous phase at −40 ms, 10.7 Hz for the frontal ROI. Phosphene perception oscillates along with the alpha phase (two-way repeated-measures ANOVA: F (1,8) = 3.360, p-value = 0.0028 eta^2^ = 29.58), with an optimal phase for phosphene perception at the phase π/2 of the alpha cycle. The percentage of variance explained by the phase is more important for high-(21.2% difference between π/2 and − π/2) compared to low-(17.6%) alpha amplitude trials.

We repeated this analysis for the individual electrodes PO3 and AFz and observed similar effects. A two-way repeated-measures ANOVA showed a significant effect of the phase for both electrodes PO3 (F(1,8) = 2.113, p-value = 0.0472, eta^2^ = 20.89, SS = 0.937) and AFz (F(1,8) = 2.106, p-value = 0.0479, eta^2^ = 20.84, SS = 0.908), no significant main effect of the amplitude for either electrode PO3 (F(1,8) = 0.461, p-value = 0.5164, eta^2^ = 5.45, SS = 0.002) or AFz (F(1,8) = 0.002, p-value = 0.9633, eta^2^ = 0.03, SS < 0.001), and no interaction for either electrode PO3 (F(1,8) = 0.612, p-value = 0.7648, SS = 0.294) or AFz (F(1,8) = 0.389, p-value = 0.9224, SS = 0.2).

To understand the role of spontaneous alpha oscillations phase-amplitude tradeoffs on cortical excitability and subsequent perceptual performance, we then focused on the post-pulse evoked activity. Dugué et al. (2011a) previously observed a larger post-pulse ERP in the perceived-than in the unperceived-phosphene trials, with a positive differential activity for PO3, and negative for AFz, between ~300 ms and ~600 ms. They interpreted these results as a physiological consequence of phosphene perception. Here, we computed the ERP difference between perceived- and unperceived-phosphene conditions, separately for low- and high-alpha amplitude trials and observed a similar effect in both low- and high-alpha amplitude conditions (**Figure 3**; see also **Extended Data Figure 3-1** for ERPs on each condition separately). An FFT was computed from 400 to 800 ms after the pulse on the perceived- and unperceived phosphene ERP, separately for electrodes PO3 and AFz, and for low- and high-alpha amplitude trials. There was no significant frequency peak at 10.24 Hz in the post-pulse ERP amplitude spectra, in any of the conditions for both electrode PO3 (one-tailed t-tests against the 1/f aperiodic component: low-alpha amplitude, perceived: p-value = 0.9244, Cohen’s d = −0.4967, CI = [−28.6420; Infinity]; unperceived: p-value = 0.6837, Cohen’s d = −0.1176, CI = [−11.6204; Infinity]; high-alpha amplitude, perceived: p-value = 0.4279, Cohen’s d = 0.0643, CI = [−17.1197; Infinity]; unperceived: p-value = 0.0971, Cohen’s d = 0.3345, CI = [−4.1354; Infinity]) and AFz (low-alpha amplitude, perceived: 0.9864, Cohen’s d = −1.3376, CI = [−36.8469; Infinity]; unperceived: 0.5615, Cohen’s d = −0.0520, CI = [−15.0938; Infinity]; high-alpha amplitude, perceived: p-value = 0.3038, Cohen’s d = 0.0322, CI = [−28.0728; Infinity]; unperceived: p-value = 0.1811, Cohen’s d = 0.0610, CI = [−17.3280; Infinity]). Thus, the post-pulse signal likely does not contain sufficient pre-pulse information to translate into a contamination of the post-pulse ERP.

Finally, we investigated the link between the pre-pulse alpha phase and amplitude, and the post-pulse evoked activity. For each participant, we sorted the low- and high-alpha amplitude trials in 9 bins, as previously described, separately for electrodes PO3 and AFz. For each phase bin and alpha-amplitude condition, the maximum perceived-unperceived ERP difference was computed on a selected time-window of interest (see **Materials & Methods**; gray shaded areas in **Figure 3**). A two-way repeated-measures ANOVA on electrode PO3 (**Figure 6A**) revealed a significant main effect of the phase (F(1,8) = 2.338, p-value = 0.0286, eta^2^ = 22.62, SS = 430.66) and alpha-amplitude (F(1,8) = 13.623, p-value = 0.0061, eta^2^ = 63, SS = 137.62; this is coherent with the selection of the ERP time window of interest and will not be further interpreted; see **Materials & Methods**), but no significant interaction (F(1,8) = 0.377, p-value = 0.9289, SS = 61.76). The two-way repeated-measures ANOVA on electrode AFz (**Figure 6C**) showed a significant main effect of the alpha-amplitude (F(1,8) = 12.749, p-value = 0.0073, eta^2^ = 61.44, SS = 102.97; this is coherent with the selection of the ERP time window of interest and will not be further interpreted; see **Materials & Methods**), no significant effect of the phase (F(1,8) = 0.393, p-value = 0.9206, eta^2^ = 4.68, SS = 161.37), and no interaction (F(1,8) = 0.376, p-value = 0.9297, SS = 158.75). In other words, for both low- and high-alpha amplitude, the phase of pre-pulse spontaneous alpha oscillations predicts the ERP difference exclusively for the occipital electrode PO3. Specifically, we observed a higher ERP difference at −π/4, i.e., around the non-optimal phase for phosphene perception (see **Figure 5A**). This effect seems to come from an increased ERP in perceived-phosphene trials specifically. Indeed, we extracted the peak of the ERP for low- and high-alpha amplitude trials, at −π/4 and π/2 phases of the alpha cycle, corresponding respectively to the maximum and the minimum ERP difference observed (see **Figure 5A**), separately for perceived- and unperceived-ERPs, for the 9 participants (**Figure 6B**). We implemented two linear mixed effects models, one for each alpha-amplitude condition. In each model, we entered as fixed effects the phase (-π/4, π/2), the phosphene condition (perceived, unperceived), as well as their interaction. As random effect, we had participants’ intercepts and slopes for the effect of phase and phosphene condition. We observed a significant effect of the phosphene condition for both low-(t(32) = −3.1, p-value = 0.004, estimate = −12.201 ± 3.935, standard error) and high-(t(32) = −3.252, p-value = 0.003, estimate = −12.188 ± 3.748, standard error) alpha amplitude conditions, a significant effect of the phase for low-(t(32) = −2.551, p-value = 0.0157, estimate = −10.161 ± 3.983, standard error) and high-(t(32) = −2.313, p-value = 0.0273, estimate = −8.833 ± 3.82, standard error) alpha amplitude conditions, and a significant interaction between the phosphene condition and the phase for both low-(t(32) = 2.468, p-value = 0.0191, estimate = 6.142 ± 2.489, standard error) and high-(t(32) = 2.237, p-value = 0.033, estimate = 5.292 ± 2.369, standard error) alpha amplitude conditions. A post-hoc analysis showed that the ERP difference at −π/4 for perceived-phosphene trials was significantly higher compared to unperceived-phosphene trials at −π/4 for both low-(one-tailed t-test, p-value = 0.0265, Cohen’s d = 0.809, CI = [1.09; Infinity]) and high-(p-value = 0.0074, Cohen’s d = 1.069, CI = [2.753; Infinity]) alpha amplitude conditions, and compared to unperceived-phosphene trials at π/2 for both low-(p-value = 0.0254, Cohen’s d = 0.477, CI = [0.747; Infinity]) and high-(p-value = 0.0253, Cohen’s d = 0.737, CI = [0.984; Infinity]) alpha amplitude conditions. Thus, around −π/4 for both low- and high-alpha amplitude, we observed a low percentage of perceived-phosphene (**Figure 5A**) associated with a high ERP difference when the phosphene is perceived (**Figure 6A**).

**Figure 6.**
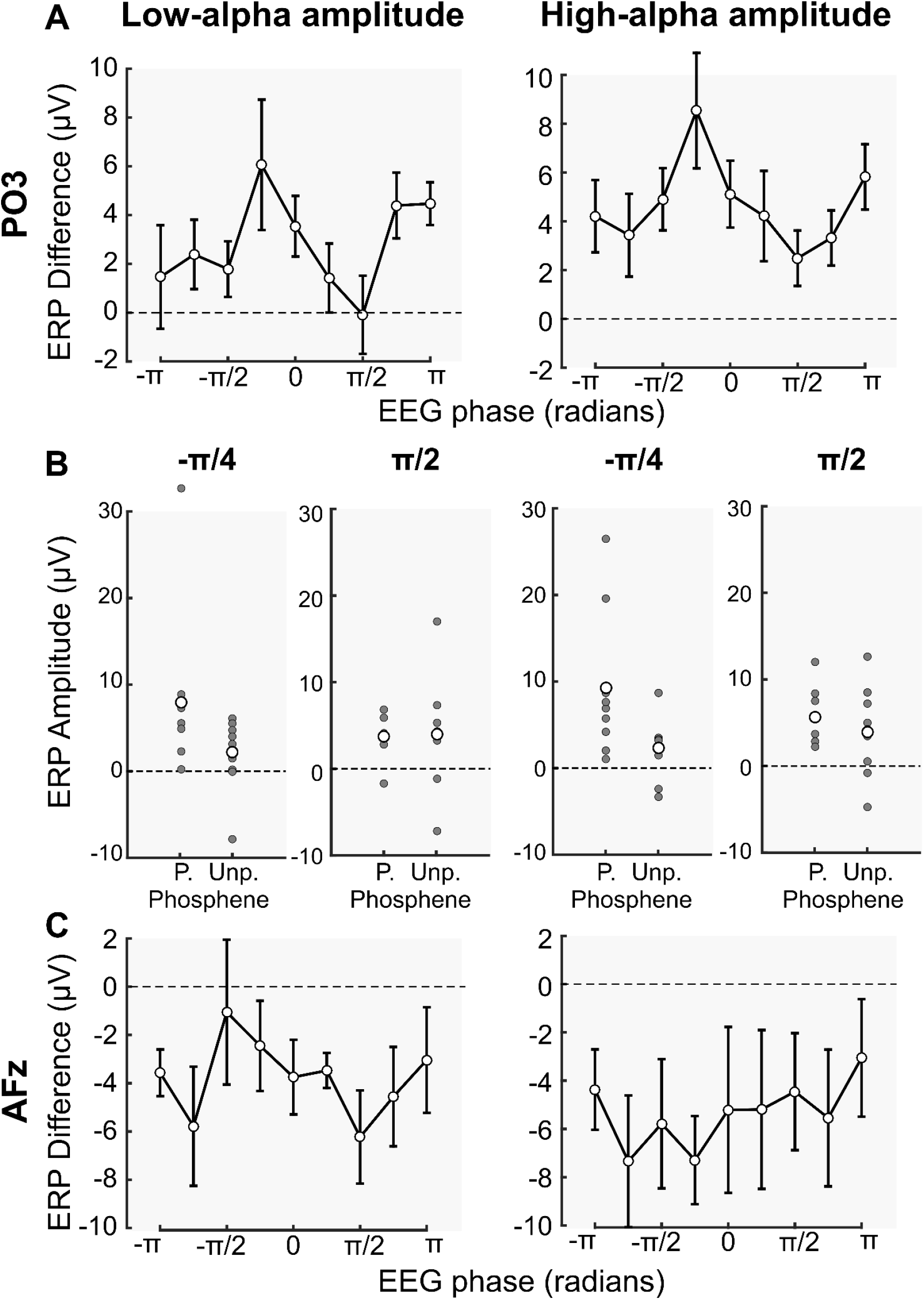
The ERP is higher for perceived-phosphene trials at the non-optimal phase for phosphene perception, for the electrode PO3. Left panels, ERP difference between perceived- and unperceived-phosphene conditions computed for 9 phase bins, and averaged across the 9 participants, for low-alpha amplitude trials. Right panels, for high-alpha amplitude trials. **A.** Single trials were sorted into 9 phase bins according to the instantaneous phase at −77 ms, 10.7 Hz, for the electrode PO3. For each phase bin, the maximum perceived-unperceived ERP difference was computed. Errors bars, standard error of the mean. The ERP difference oscillates along with the alpha phase (F(1,8) = 2.338, p-value = 0.0286, eta^2^ = 22.62). The ERP difference was higher at the phase −π/4. **B.** ERP for the perceived- and unperceived-phosphene conditions, for the electrode PO3, for the phase −π/4 and π/2. P., perceived-; Unp., unperceived-phosphene conditions. Gray dots, maximum ERP for each participant. White dots, averaged ERP across the 9 participants. The maximum ERP at the phase −π/4 for perceived-phosphene trials was significantly higher compared to unperceived-phosphene trials at the phase −π/4 for both low-(one-tailed t-test, p-value = 0.0265) and high-(p-value = 0.0074) amplitude trials-type, and compared to unperceived-phosphene trials at the phase π/2 for both low-(p-value = 0.0254) and high-(p-value = 0.0253) amplitude trials-type. **C.** Single trials were sorted into 9 phase bins according to the instantaneous phase at −40 ms, 10.7 Hz, for the electrode AFz. For each phase bin, the maximum perceived-unperceived ERP difference was computed. Errors bars, standard error of the mean.

## Discussion

In this study, we tested the two clear predictions of the Pulsed Inhibition theory (Jensen and Mazaheri, 2010; Klimesch et al., 2007; Mathewson et al., 2011): (1) high alpha amplitude induces cortical inhibition at specific phases of the alpha cycle, leading to periodic perceptual performance; while (2) low alpha amplitude is less susceptible to phasic (periodic) pulsed inhibition, leading to overall higher perceptual performance. Cortical excitability was assessed by both phosphene detection and post-pulse evoked EEG activity. We showed that the pre-pulse phase of spontaneous alpha oscillations (~10 Hz) modulates the probability to perceive a phosphene (with a non-optimal phase between −π/2 and −π/4). This phase effect was stronger for high-alpha amplitude trials. Moreover, the pre-pulse non-optimal phase leads to an increase in post-pulse evoked activity (ERP), in phosphene-perceived trials specifically. Together, our results provide strong evidence in favor of the Pulsed Inhibition theory by establishing a causal link between the amplitude and the phase of spontaneous alpha oscillations, cortical excitability, and subsequent perceptual performance.

### Alpha phase-amplitude tradeoffs on perception

As previously described in the literature, we found that the phase of spontaneous oscillations in the alpha frequency range predicts whether a near-threshold stimulus would be successfully perceived (Busch et al., 2009; Dugué et al., 2011a; Mathewson et al., 2009; Samaha et al., 2015, 2017). The use of TMS to induce phosphene perception rather than an external stimulation allows for direct access to the absolute phase of spontaneous oscillations. We found that a phase between −π/2 and −π/4 was associated with inhibitory moments leading to lower perceptual performance while the opposite one (π/2) was optimal for perception. Critically, as predicted by the Pulsed Inhibition theory, our results are in line with some previous studies showing that the phase of spontaneous alpha oscillations better predicts perceptual performance for high than for low-alpha amplitude (Ai and Ro, 2014; Alexander et al., 2020; Bonnefond and Jensen, 2015; Herrmann et al., 2016; Kizuk and Mathewson, 2017; Mathewson et al., 2009; Ng et al., 2012; Spitzer et al., 2016) but not others (Busch and VanRullen, 2010; Harris et al., 2018; Milton and Pleydell-Pearce, 2016; Zoefel and Heil, 2013; Madsen et al., 2019). Other studies investigated the specific case in which a high-alpha amplitude condition is compared to the actual absence of alpha oscillations (Bergmann et al., 2019; Schaworonkow et al., 2019, 2018; Stefanou et al., 2018; Zrenner et al., 2018). They found periodic functional inhibition induced by mu oscillations (alpha oscillations recorded in the motor cortex) in the high-alpha amplitude condition (see next paragraph for more details). Here, we compared high-alpha amplitude trials to trials in which alpha oscillations were present but with a lower amplitude, and found a phase effect in both alpha-amplitude conditions, but strongest when alpha amplitude is high.

### Alpha phase-amplitude tradeoffs on cortical excitability

We observed that the phase and the amplitude of spontaneous alpha oscillations influence cortical excitability, only when there is subsequent perception. Indeed, a phase between −π/2 and −π/4 led to higher ERP exclusively for phosphene perception trials. Interestingly, this phase was also associated with lower perceptual performance. The non-optimal phase of the alpha oscillations (between −π/2 and −π/4) tends to create periodic inhibitory cortical states favoring the absence of phosphene perception, which leads to a greater ERP response when a phosphene is in fact perceived.

It is important to notice that the paradigm developed by Dugué et al. (2011a) was designed to specifically investigate the role of the phase of alpha oscillations (and not the phase-amplitude tradeoffs). Yet, the results are compelling and in line with other studies observing similar effects (Bonnefond and Jensen, 2015; Hussain et al., 2019; and others comparing high-alpha amplitude to the absence of alpha oscillations: Bergmann et al., 2019; Schaworonkow et al., 2019, 2018; Stefanou et al., 2018; Zrenner et al., 2018). In the motor modality, they used single-pulse TMS over the motor cortex to induce motor evoked potentials (MEP) allowing to estimate corticospinal excitability. They found an increase in corticospinal excitability and the subsequent MEP for high-mu amplitude oscillations (i.e., alpha oscillations observed in somatosensory and motor areas) and for specific mu phases (Bergmann et al., 2019; Hussain et al., 2019; Schaworonkow et al., 2019, 2018; Stefanou et al., 2018; Zrenner et al., 2018). Bonnefond and Jensen (2015) alternatively analyzed the power of high gamma oscillations (80-120 Hz) considered to reflects neuronal firing (Ray et al., 2008). They showed that gamma power was weaker at the trough of high-alpha amplitude oscillations (Bonnefond and Jensen, 2015). As predicted by the Pulsed Inhibition theory, high-alpha amplitude modulates cortical and corticospinal excitability periodically.

Interestingly, although the Pulsed Inhibition theory (Jensen and Mazaheri, 2010) originally proposed asymmetrical pulsed inhibition (i.e., inhibition at one particular phase and no inhibition at the opposite one), Bergmann et al. (2019) argued in favor of asymmetrical pulsed facilitation. Indeed, they assessed the role of the GABAergic system, considered the main source of inhibition in the brain (Ribak and Yan, 2000), on the amplitude and phase of mu oscillations, and did not observe any relation. The symmetry/asymmetry hypothesis was not explicitly assessed in the present study. Further investigation is thus necessary to disentangle the three possibilities: (1) symmetric pulsed inhibition and facilitation, (2) asymmetrical pulsed inhibition or (3) asymmetrical pulsed facilitation.

### Alpha, a top-down process?

Our results show a potential functional link between the occipital and the frontal lobes. Several authors have proposed that alpha carries feedback information (Michalareas et al., 2016; van Kerkoerle et al., 2014) and that the amplitude of occipital alpha oscillations is modulated by top-down connections from fronto-parietal regions (Klimesch et al., 2007; Mathewson et al., 2011). Here, we can speculate that the frontal region plays a role in the emergence of inhibitory and excitatory moments in occipital cortex. Specifically, their top-down influence on the amplitude of alpha oscillations would enhance or reduce locally the effect of the phase of occipital alpha oscillations on perceptual performance, thus explaining that the link between the phase and the amplitude of alpha oscillations and cortical excitability (ERP) was only present in the occipital ROI. In addition, the previous study from which the data originate (Dugué et al, 2011) shows that the time at which the phase predicted the perceptual outcome differed by nearly one-half alpha-cycle between the occipital (−77 ms) and the frontal (−40 ms) ROI. This difference may reflect the delay for neural information to be transferred from one brain region to the other, consistent with previous observations of an alpha phase difference between occipital and frontal regions during visual perception (Burkitt et al., 2000; Patten et al., 2012; Alamia and VanRullen, 2019; Pang et al., 2020; Tsoneva et al., 2021). Further studies are warranted to investigate the functional interplay between the frontal and occipital cortex in the context of the Pulsed Inhibition theory.

### Conclusion

Our study provides strong causal evidence in favor of tradeoffs between the phase and the amplitude of alpha oscillations to create periodic inhibitory moments leading to rhythms in perception. As predicted by the Pulsed Inhibition theory, the effect of the phase of spontaneous alpha oscillations on perception increases for larger alpha amplitude.

## References

Ai L, Ro T (2014) The phase of prestimulus alpha oscillations affects tactile perception. J Neurophysiol 111:1300–1307.

Alamia A, VanRullen R (2019) Alpha oscillations and traveling waves: Signatures of predictive coding?. PLOS Biology 17(10): e3000487: 1–26.

Alexander KE, Estepp JR, Elbasiouny SM (2020) Effects of Neuronic Shutter Observed in the EEG Alpha Rhythm. eNeuro 7:1–14.

Bergmann TO, Lieb A, Zrenner C, Ziemann U (2019) Pulsed Facilitation of Corticospinal Excitability by the Sensorimotor μ-Alpha Rhythm. J Neurosci Off J Soc Neurosci 39:10034–10043.

Bollimunta A, Chen Y, Schroeder CE, Ding M (2008) Neuronal mechanisms of cortical alpha oscillations in awake-behaving macaques. J Neurosci Off J Soc Neurosci 28:9976–9988.

Bonnefond M, Jensen O (2015) Gamma activity coupled to alpha phase as a mechanism for top-down controlled gating. PloS One 10:e0128667:1–11.

Burkitt, G. R., Silberstein, R. B., Cadusch, P. J., & Wood, A. W. (2000) Steady-state visual evoked potentials and travelling waves. Clinical Neurophysiology, 111(2), 246–258.

Busch NA, Dubois J, VanRullen R (2009) The phase of ongoing EEG oscillations predicts visual perception. J Neurosci Off J Soc Neurosci 29:7869–7876.

Busch NA, VanRullen R (2010) Spontaneous EEG oscillations reveal periodic sampling of visual attention. Proc Natl Acad Sci U S A 107:16048–16053.

Clayton MS, Yeung N, Cohen Kadosh R (2018) The many characters of visual alpha oscillations. Eur J Neurosci 48:2498–2508.

Dugué L, Beck A-A, Marque P, VanRullen R (2019) Contribution of FEF to Attentional Periodicity during Visual Search: A TMS Study. eNeuro 6:1–10.

Dugué L, Marque P, VanRullen R (2015) Theta oscillations modulate attentional search performance periodically. J Cogn Neurosci 27:945–958.

Dugué L, Marque P, VanRullen R (2011a) The phase of ongoing oscillations mediates the causal relation between brain excitation and visual perception. J Neurosci Off J Soc Neurosci 31:11889–11893.

Dugué L, Marque P, VanRullen R (2011b) Transcranial magnetic stimulation reveals attentional feedback to area V1 during serial visual search. PloS One 6:e19712:1–8.

Dugué L, Roberts M, Carrasco M (2016) Attention Reorients Periodically. Curr Biol 26:1595–1601.

Dugué L, VanRullen R (2017) Transcranial Magnetic Stimulation Reveals Intrinsic Perceptual and Attentional Rhythms. Front Neurosci 11:1–7.

Ergenoglu T, Demiralp T, Bayraktaroglu Z, Ergen M, Beydagi H, Uresin Y (2004) Alpha rhythm of the EEG modulates visual detection performance in humans. Brain Res Cogn Brain Res 20:376–383.

Gerwig, Marcus, Ludwig Niehaus, Oliver Kastrup, Philipp Stude, et Hans C. Diener (2005) Visual Cortex Excitability in Migraine Evaluated by Single and Paired Magnetic Stimuli. Headache: The Journal of Head and Face Pain 45: 1394‑99.

Goldman RI, Stern JM, Engel J, Cohen MS (2002) Simultaneous EEG and fMRI of the alpha rhythm. Neuroreport 13:2487–2492.

Haegens S, Barczak A, Musacchia G, Lipton ML, Mehta AD, Lakatos P, Schroeder CE (2015) Laminar Profile and Physiology of the α Rhythm in Primary Visual, Auditory, and Somatosensory Regions of Neocortex. J Neurosci Off J Soc Neurosci 35:14341–14352.

Haegens S, Nácher V, Luna R, Romo R, Jensen O (2011) α-Oscillations in the monkey sensorimotor network influence discrimination performance by rhythmical inhibition of neuronal spiking. Proc Natl Acad Sci U S A 108:19377–19382.

Händel BF, Haarmeier T, Jensen O (2011) Alpha oscillations correlate with the successful inhibition of unattended stimuli. J Cogn Neurosci 23:2494–2502.

Harris AM, Dux PE, Mattingley JB (2018) Detecting Unattended Stimuli Depends on the Phase of Prestimulus Neural Oscillations. J Neurosci Off J Soc Neurosci 38:3092–3101.

Herrmann B, Henry MJ, Haegens S, Obleser J (2016) Temporal expectations and neural amplitude fluctuations in auditory cortex interactively influence perception. NeuroImage 124:487–497.

Hussain SJ, Claudino L, Bönstrup M, Norato G, Cruciani G, Thompson R, Zrenner C, Ziemann U, Buch E, Cohen LG (2019) Sensorimotor Oscillatory Phase-Power Interaction Gates Resting Human Corticospinal Output. Cereb Cortex N Y N 1991 29:3766–3777.

Jensen O, Mazaheri A (2010) Shaping functional architecture by oscillatory alpha activity: gating by inhibition. Front Hum Neurosci 4:1–13.

Kammer, Thomas, et Lisa W. Baumann (2010) Phosphene Thresholds Evoked with Single and Double TMS Pulses. Clinical Neurophysiology 121: 376‑79.

Kienitz, R., M. C. Schmid, et L. Dugué (2021) Rhythmic Sampling Revisited: Experimental Paradigms and Neural Mechanisms. Eur J Neurosci. 1–15.

Kizuk SAD, Mathewson KE (2017) Power and Phase of Alpha Oscillations Reveal an Interaction between Spatial and Temporal Visual Attention. J Cogn Neurosci 29:480–494.

Klimesch W, Sauseng P, Hanslmayr S (2007) EEG alpha oscillations: the inhibition-timing hypothesis. Brain Res Rev 53:63–88.

Lin Y-J, Shukla L, Dugué L, Valero-Cabré A, Carrasco M (2021) Transcranial magnetic stimulation entrains alpha oscillatory activity in occipital cortex. Sci Rep 11:18562.

Madsen KH, Karabanov AN, Krohne LG, Safeldt MG, Tomasevic L, Siebner HR (2019) No trace of phase: Corticomotor excitability is not tuned by phase of pericentral mu-rhythm. Brain Stimulat 12:1261–1270.

Mathewson KE, Gratton G, Fabiani M, Beck DM, Ro T (2009) To see or not to see: prestimulus alpha phase predicts visual awareness. J Neurosci Off J Soc Neurosci 29:2725–2732.

Mathewson KE, Lleras A, Beck DM, Fabiani M, Ro T, Gratton G (2011) Pulsed out of awareness: EEG alpha oscillations represent a pulsed-inhibition of ongoing cortical processing. Front Psychol 2:1–15.

Michalareas G, Vezoli J, van Pelt S, Schoffelen J-M, Kennedy H, Fries P (2016) Alpha-Beta and Gamma Rhythms Subserve Feedback and Feedforward Influences among Human Visual Cortical Areas. Neuron 89:384–397.

Milton A, Pleydell-Pearce CW (2016) The phase of pre-stimulus alpha oscillations influences the visual perception of stimulus timing. NeuroImage 133:53–61.

Moosmann M, Ritter P, Krastel I, Brink A, Thees S, Blankenburg F, Taskin B, Obrig H, Villringer A (2003) Correlates of alpha rhythm in functional magnetic resonance imaging and near infrared spectroscopy. NeuroImage 20:145–158.

Ng BSW, Schroeder T, Kayser C (2012) A precluding but not ensuring role of entrained low-frequency oscillations for auditory perception. J Neurosci Off J Soc Neurosci 32:12268–12276.

Pang, Z., Alamia, A., & VanRullen, R. (2020) Turning the Stimulus On and Off Changes the Direction of α Traveling Waves. Eneuro, 7(6): 1–11.

Patten TM, Rennie CJ, Robinson PA, Gong P (2012) Human Cortical Traveling Waves: Dynamical Properties and Correlations with Responses. PLOS ONE 7(6): e38392: 1–10.

Pfurtscheller G, Lopes da Silva FH (1999) Event-related EEG/MEG synchronization and desynchronization: basic principles. Clin Neurophysiol 110:1842–1857.

Ray, Patty G., Kimford J. Meador, Charles M. Epstein, David W. Loring, et Larry J. Day (1998) Magnetic Stimulation of Visual Cortex: Factors Influencing the Perception of Phosphenes. Journal of Clinical Neurophysiology 15: 351–357.

Ray S, Crone NE, Niebur E, Franaszczuk PJ, Hsiao SS (2008) Neural Correlates of High-Gamma Oscillations (60-200 Hz) in Macaque Local Field Potentials and Their Potential Implications in Electrocorticography. J Neurosci 28:11526–11536.

Ribak CE, Yan X-X (2000) GABA neurons in the neocortex. In: Martin DL, Olsen RW, editors. GABA in the nervous system: the view at fifty years. Philadelphia: Lippincott Williams and Wilkins. p 357–68.

Romei V, Brodbeck V, Michel C, Amedi A, Pascual-Leone A, Thut G (2008) Spontaneous fluctuations in posterior alpha-band EEG activity reflect variability in excitability of human visual areas. Cereb Cortex N Y N 1991 18:2010–2018.

Rossi S, Hallett M, Rossini PM, Pascual-Leone A, Safety of TMS Consensus Group (2009) Safety, ethical considerations, and application guidelines for the use of transcranial magnetic stimulation in clinical practice and research. Clin Neurophysiol Off J Int Fed Clin Neurophysiol 120:2008–2039.

Samaha J, Bauer P, Cimaroli S, Postle BR (2015) Top-down control of the phase of alpha-band oscillations as a mechanism for temporal prediction. Proc Natl Acad Sci U S A 112:8439–8444.

Samaha J, Iemi L, Haegens S, Busch NA (2020) Spontaneous Brain Oscillations and Perceptual Decision-Making. Trends Cogn Sci 24:639–653.

Samaha J, Gosseries O, Postle BR (2017) Distinct Oscillatory Frequencies Underlie Excitability of Human Occipital and Parietal Cortex. J Neurosci 37:2824–2833.

Sauseng P, Klimesch W, Stadler W, Schabus M, Doppelmayr M, Hanslmayr S, Gruber WR, Birbaumer N (2005) A shift of visual spatial attention is selectively associated with human EEG alpha activity. Eur J Neurosci 22:2917–2926.

Schaworonkow N, Caldana Gordon P, Belardinelli P, Ziemann U, Bergmann TO, Zrenner C (2018) μ-Rhythm Extracted With Personalized EEG Filters Correlates With Corticospinal Excitability in Real-Time Phase-Triggered EEG-TMS. Front Neurosci 12:1–6.

Schaworonkow N, Triesch J, Ziemann U, Zrenner C (2019) EEG-triggered TMS reveals stronger brain state-dependent modulation of motor evoked potentials at weaker stimulation intensities. Brain Stimulat 12:110–118.

Scheeringa R, Mazaheri A, Bojak I, Norris DG, Kleinschmidt A (2011) Modulation of visually evoked cortical FMRI responses by phase of ongoing occipital alpha oscillations. J Neurosci Off J Soc Neurosci 31:3813–3820.

Spitzer B, Blankenburg F, Summerfield C (2016) Rhythmic gain control during supramodal integration of approximate number. NeuroImage 129:470–479.

Stefanou M-I, Desideri D, Belardinelli P, Zrenner C, Ziemann U (2018) Phase Synchronicity of μ-Rhythm Determines Efficacy of Interhemispheric Communication Between Human Motor Cortices. J Neurosci Off J Soc Neurosci 38:10525–10534.

Taylor PCJ, Walsh V, Eimer M (2010) The neural signature of phosphene perception. Hum Brain Mapp 31:1408–1417.

Thut G, Nietzel A, Brandt SA, Pascual-Leone A (2006) Alpha-band electroencephalographic activity over occipital cortex indexes visuospatial attention bias and predicts visual target detection. J Neurosci Off J Soc Neurosci 26:9494–9502.

Tsoneva, T., Garcia-Molina, G., & Desain, P. (2021) SSVEP phase synchronies and propagation during repetitive visual stimulation at high frequencies. Scientific reports, 11(1), 1–13.

van Dijk H, Schoffelen J-M, Oostenveld R, Jensen O (2008) Prestimulus oscillatory activity in the alpha band predicts visual discrimination ability. J Neurosci Off J Soc Neurosci 28:1816–1823.

van Kerkoerle T, Self MW, Dagnino B, Gariel-Mathis M-A, Poort J, van der Togt C, Roelfsema PR (2014) Alpha and gamma oscillations characterize feedback and feedforward processing in monkey visual cortex. Proc Natl Acad Sci U S A 111:14332–14341.

VanRullen R (2016a) Perceptual Cycles. Trends Cogn Sci 20:723–735.

VanRullen R (2016b) How to Evaluate Phase Differences between Trial Groups in Ongoing Electrophysiological Signals. Front Neurosci 10:1−22.

Varela FJ, Toro A, John ER, Schwartz EL (1981) Perceptual framing and cortical alpha rhythm. Neuropsychologia 19:675–686.

Zoefel B, Heil P (2013) Detection of Near-Threshold Sounds is Independent of EEG Phase in Common Frequency Bands. Front Psychol 4:1–17.

Zrenner C, Desideri D, Belardinelli P, Ziemann U (2018) Real-time EEG-defined excitability states determine efficacy of TMS-induced plasticity in human motor cortex. Brain Stimulat 11:374–389.

